# SEAHORSE: A Serendipity Engine Assaying Heterogeneous Omics-Related Sampling Experiments

**DOI:** 10.1101/2025.08.15.670514

**Authors:** Adam Quackenbush, Jaya Kolluri, Rohan Biju, Saron Nhong, Derrick K DeConti, Katherine H. Shutta, Helena Wu, John Quackenbush, Enakshi Saha, Tara Eicher

## Abstract

Large public molecular atlases such as the Genotype-Tissue Expression (GTEx) project and The Cancer Genome Atlas (TCGA) invite systematic discovery, yet most analyses remain hypothesis-driven and interrogate a tiny fraction of possible relationships among phenotypic, clinical, and molecular variables. We developed SEAHORSE (Serendipity Engine Assaying Heterogeneous Omics-Related Sampling Experiments), a discovery engine and accompanying R package that exhaustively precomputes all pairwise associations across heterogeneous data types and presents them as a searchable association landscape. Using GTEx (948 donors, 43 tissues, 154 phenotypes), SEAHORSE generated 341,008 phenotype–phenotype associations, 125,246,938 phenotype–gene associations, and 10,269,910,410 gene–gene correlations. In parallel analyses spanning 33 tumor types in TCGA, SEAHORSE generated 625,042 phenotype–phenotype associations, 183,080,369 phenotype–gene associations, and 12,096,948,950 gene–gene correlations. Across GTEx, height was repeatedly associated with enrichment of transcriptional programs, most strikingly the Kyoto Encyclopedia of Genes and Genomes (KEGG) term “Pathways in Cancer,” significant in 17 tissues, offering a molecular entry point into long-reported links between stature and cancer risk. Height was also associated with immune, cardiovascular, and neurologic programs. In TCGA, age was consistently associated with WNT signaling, translation, cell differentiation, and cell cycle programs across tumors.

These findings illustrate a new paradigm: large cohorts should be treated not merely as repositories for testing preconceived hypotheses but as association landscapes that can generate unexpected biological hypotheses.

**Graphical Abstract:** SEAHORSE ecosystem, including the statistical framework, Amazon Web Services (AWS)-hosted data for GTEx and TCGA, and portal.

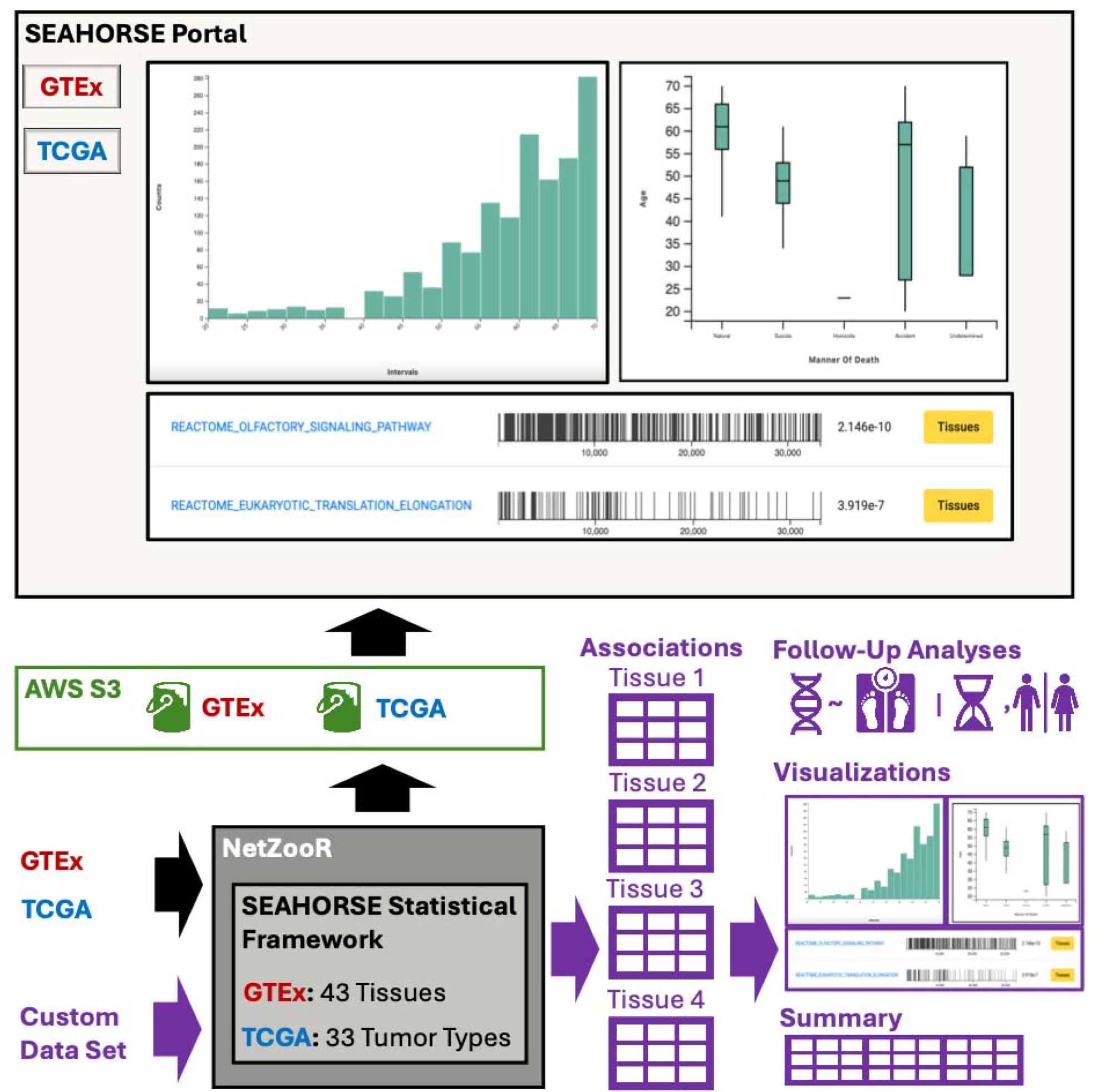

## Introduction

Modern biology has been defined not merely by the accumulation of data, but by a qualitative change in the kinds of questions those data now allow us to ask. Large cohort-based projects routinely generate molecular profiles at an unprecedented scale and are accompanied by detailed demographic, pathological, lifestyle, and clinical annotations. These resources have dramatically altered how we think about human variation, tissue-specific regulation, and disease by allowing us to test hypotheses relevant to understanding human health and disease. Among the most important early examples are the Genotype-Tissue Expression (GTEx) project, which maps transcriptional regulation across human tissues ^1 2 3^, and The Cancer Genome Atlas (TCGA), which has defined the molecular architecture of human tumors at scale ^4^. These resources offer uniquely rich environments in which diverse phenotypic and multi-omic data have been assembled in a consistent fashion for large numbers of samples.

And yet, despite the size and scope of these data sets, the majority of analyses performed on them are driven by targeted hypotheses. Investigators typically begin with a gene, pathway, or clinical parameter of interest, and then ask whether that feature differs across states or correlates with a chosen phenotype. Other studies choose related phenotypic states, such as health and disease, and compare molecular or clinical features to identify those features that distinguish them. These conventional approaches remain powerful and indispensable modes of science based on directed hypothesis testing and comparison of biological groups or subtypes. But they are, by design, conservative. They are largely based on what we already think is important, and for this reason, they risk missing relationships that lie outside our current conceptual map. The challenge is not that hypothesis-driven science is wrong; rather, it is incomplete when confronted with data spaces that are too large to navigate solely by intuition.

This tension has become increasingly explicit in discussions of data-intensive science. Several authors have argued that discovery-driven analyses can reveal biologically meaningful relationships that were not obvious from prior theory alone ^5–7^. Similarly, methods developed to identify non-linear or otherwise unexpected associations in large data sets demonstrate that the search space of plausible relationships is much larger than that most routine analyses interrogate ^8^. In biology, where multi-omic data sets now routinely contain tens of thousands of molecular features and hundreds of annotation variables, the combinatorial space of possible associations is enormous. Most of it is never examined.

We have found it helpful to think about this problem using the Johari window, a framework originally developed in psychology to classify what is known and unknown to oneself and to others—a framework now regularly used in risk assessment. Adapted to science (Figure 1), the “known knowns” correspond to well-established biology, while the “known unknowns” represent explicit gaps that can be addressed by conventional hypothesis testing. But two additional categories matter deeply for discovery: “unknown knowns,” meaning associations that may already exist in the data or literature but are not currently in the investigator’s field of view, and “unknown unknowns,” meaning relationships that are both unanticipated and uncatalogued. Large, well-annotated molecular cohorts are fertile ground for exploration of these latter categories, but only if we give ourselves tools capable of searching them.

**Figure 1.**
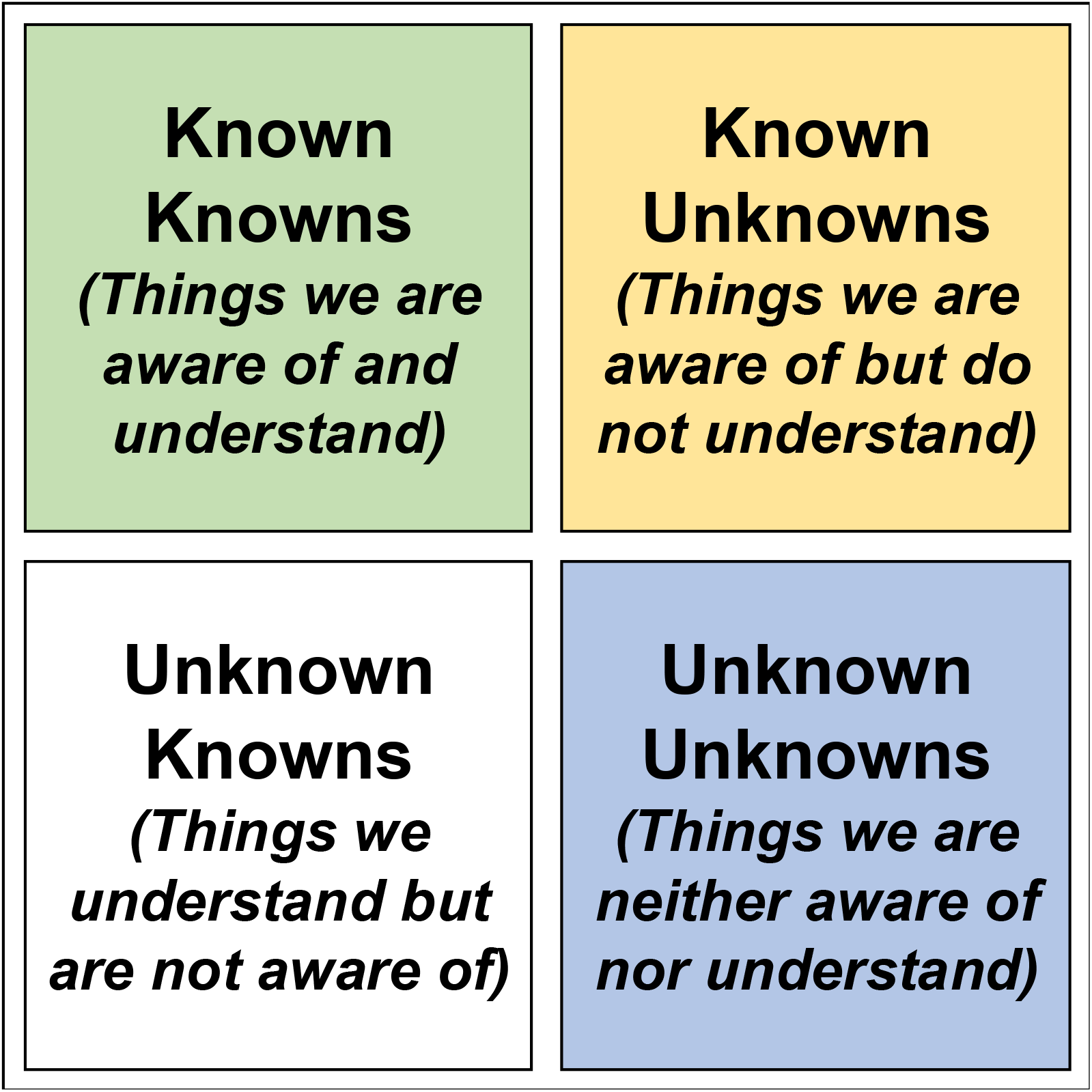
The Johari Window was first developed in psychology to classify what we and others know about ourselves. It has subsequently been generalized for use in many areas, including risk assessment, where the most dangerous elements are those we simply do not recognize as unknown to us (either because we are blind to them or because they are hidden from us). In science, the “Known Knowns” encapsulate our current understanding, and the “Known Unknowns” are the areas we explore through hypothesis testing experiments. The “Unknown Unknowns” are where new, unexpected hypotheses can be developed.

The Serendipity Engine Assaying Heterogeneous Omics-Related Sampling Experiments (SEAHORSE) was developed to make such searches practical. The resource is built on a simple idea: precompute the entire association space, organize it for efficient querying, and provide visual and statistical summaries that enable users to move from broad exploration to focused follow-up (Figure 2). In contrast to a conventional database that stores raw measurements or a portal that answers only a restricted class of targeted queries, SEAHORSE is intended to serve as a discovery engine (Figures 3, 4). It allows an investigator to ask not just whether a favorite gene correlates with a chosen trait, but what, across all available measurements, is associated with a variable of interest; which tissues show similar or divergent patterns; and what biological pathways appear repeatedly among associated genes.

**Figure 2.**
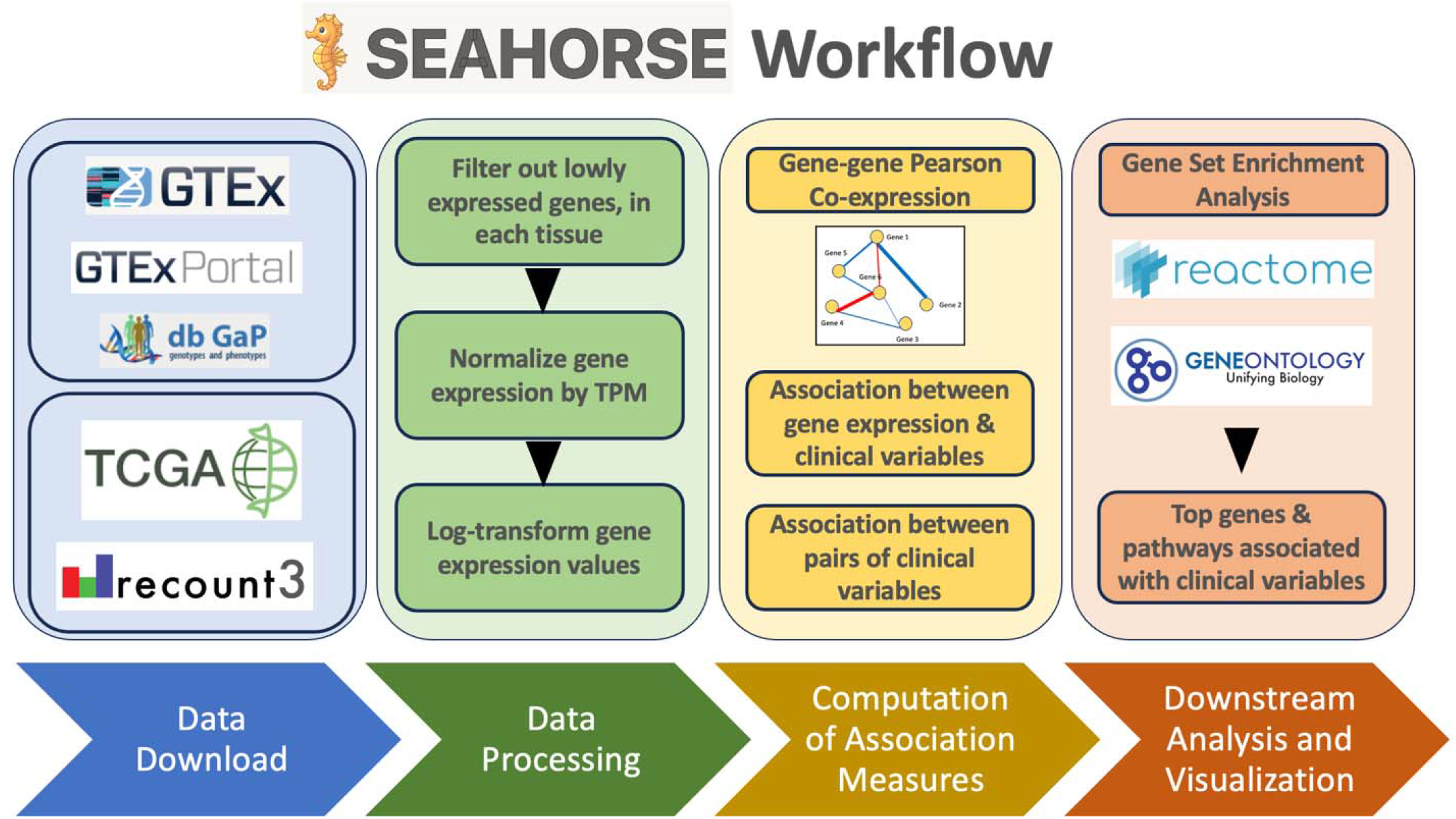
GRAPHICAL OVERVIEW of the workflow.

**Figure 3.**
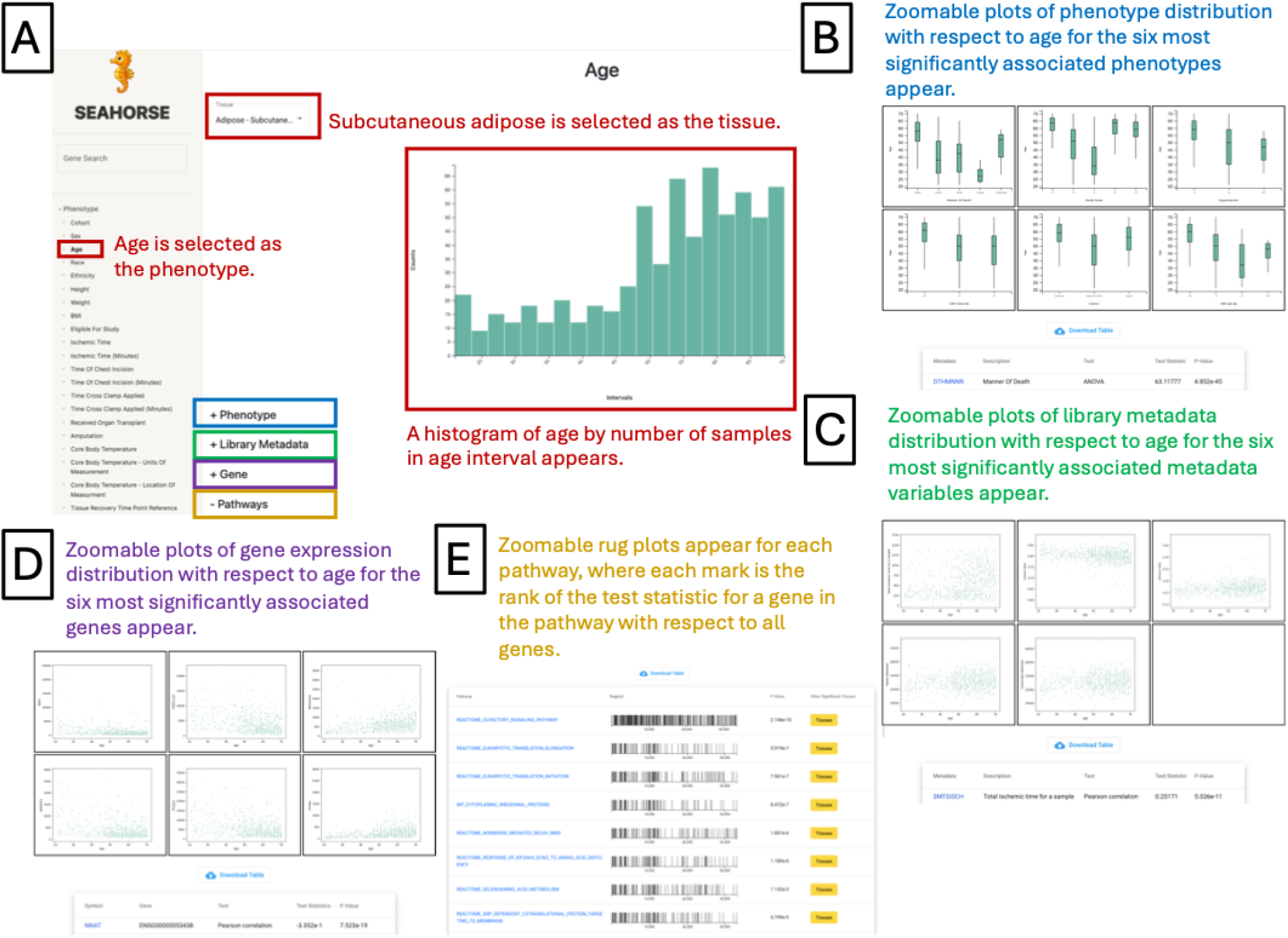
Examples of exploratory queries possible in SEAHORSE regarding phenotypes (clinical variables) in the GTEx dataset: Selecting a phenotypic variable such as “Age” and a specific tissue such as “Adipose subcutaneous” produces a display of (A) the sampling distribution of the selected phenotypic variable and four different analyses: (B) a list of other phenotypes ranked by association *p*-value of *t*-test, ANOVA, Chi-square or FFH test, or correlation test with respect to the queried phenotype, along with heat maps of the contingency table for dichotomous/nominal queried phenotypes on the *x*-axis and the other dichotomous-nominal variable on the *y*-axis, boxplots with continuous variable on the *y*-axis and dichotomous/nominal variable levels on the *x*-axis, and scatterplots with the queried continuous phenotype on the *x*-axis and the other continuous variable on the *y*-axis for the top six associations; (C) a list of library metadata variables ranked by association *p*-value with respect to the queried phenotype with the same visualizations as in B; (D) a list of genes ranked by association significance with the queried phenotype in the selected tissue along with scatterplot or boxplot with the queried phenotype and on the *x*-axis and gene expression levels on the *y*-axis for the top six associations; (E) a list of the top ten biological pathways ranked by significance of association with the selected phenotype, along with a rug plot for each pathway and a button that enables the user to list all other tissues where the selected phenotype is strongly associated with the corresponding biological pathway.

**Figure 4.**
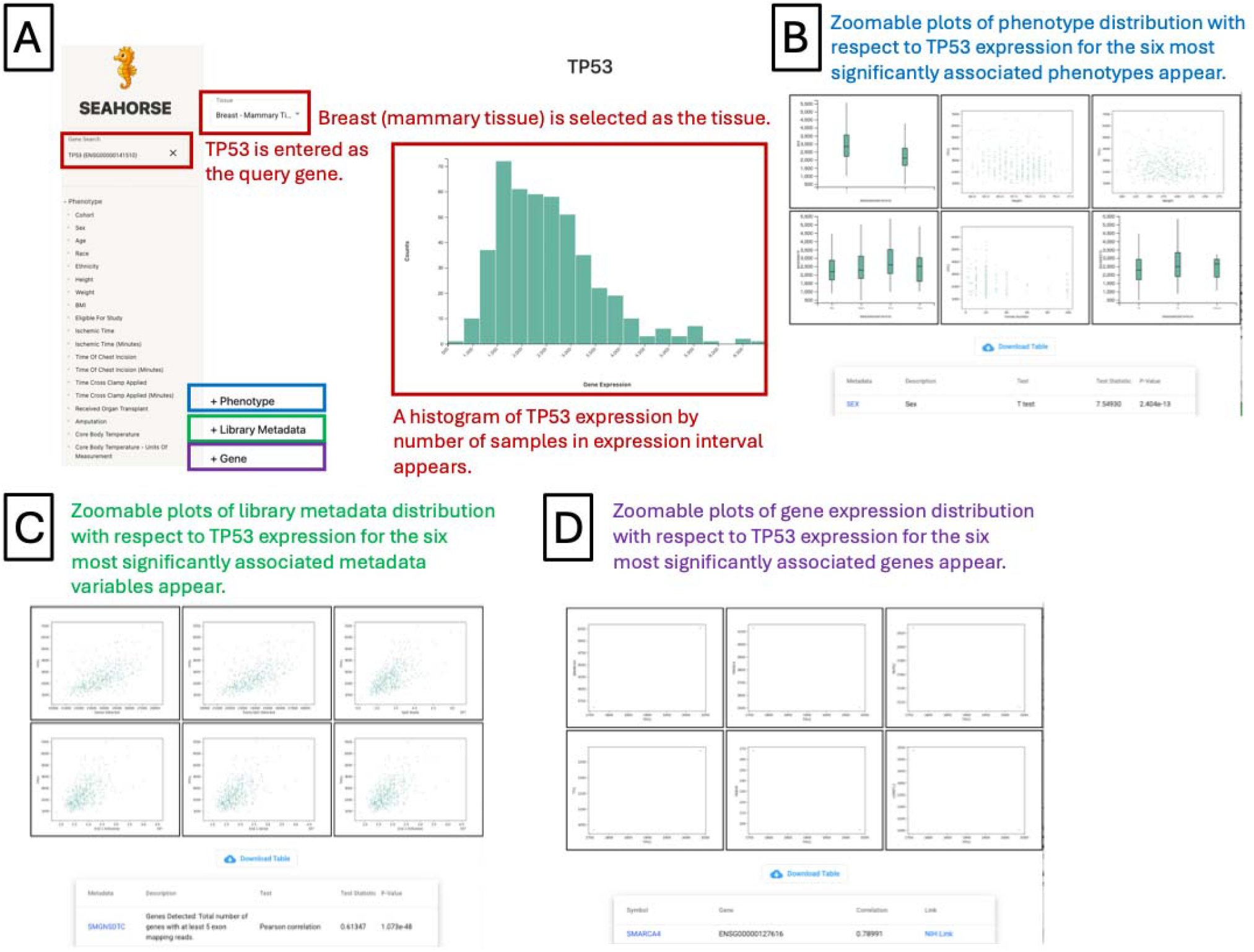
Examples of exploratory queries possible in SEAHORSE regarding specific genes in the GTEx dataset: Selecting a gene such as “TP53” and a specific tissue such as “Breast” produces a display of (A) the sampling distribution of the expression levels of the selected gene in the given tissue and three different analyses: (B) a list of phenotypes ranked by association *p*-value of t-test, ANOVA, or correlation test with respect to the queried gene, along with heat maps of the contingency table for dichotomous/nominal queried phenotypes on the *x*-axis and the other dichotomous-nominal variable on the *y*-axis, boxplots with continuous variable on the *y*-axis and dichotomous/nominal variable levels on the *x*-axis, and scatterplots with the queried continuous phenotype on the *x*-axis and the other continuous variable on the *y*-axis for the top six associations; (C) a list of library metadata variables ranked by association *p*-value with respect to the queried gene with the same visualizations as in B; (D) a list of genes ranked by correlation between the queried gene in the selected tissue along with scatterplots including the queried gene on the *x*-axis and gene expression level of the correlated gene on the *y*-axis for the top six correlations.

We chose GTEx and TCGA as the initial implementation domains for several reasons. First, both are canonical examples of modern large-scale, publicly accessible molecular cohorts. Second, both pair expression profiles with extensive metadata, making them ideal for heterogeneous all-by-all analyses. Third, both have already supported numerous important hypothesis-driven studies, allowing us to assess whether systematic exploratory analysis can recover expected biology while also uncovering less obvious relationships. In this manuscript, we describe the design of SEAHORSE, the statistical framework used to generate its precomputed association catalog (SEAHORSE statistical framework), the user-facing system used to interrogate those results (SEAHORSE portal), and a set of biological examples illustrating the kinds of unexpected observations such a platform can reveal.

Unlike query portals that primarily support targeted exploration of molecular alterations conditioned on user-selected features^1, 9–11^, SEAHORSE systematically indexes the complete heterogeneous association/correlation space (phenotype–phenotype, phenotype–gene, gene–gene) using test families matched to variable type and exposes cross-tissue recurrence and pathway-level organization as first-class outputs. The central contribution is the discovery workflow enabled by full precomputation and structured recurrence analysis, not the interface alone—moreover, the SEAHORSE statistical framework is implemented in the R package *netZooR* to allow user-driven discovery on individual data sets. While other computational tools exist to find phenotype–phenotype, phenotype–gene, and/or gene–gene associations^12–14^, SEAHORSE is unique in that it combines these functionalities on a comprehensive scale, supporting continuous, nominal, and dichotomous phenotypes, as well as both phenotype-dependent pathway enrichment analysis and follow-up phenotype–gene association analyses with adjustment for covariates (Table 1).

**Table 1.**
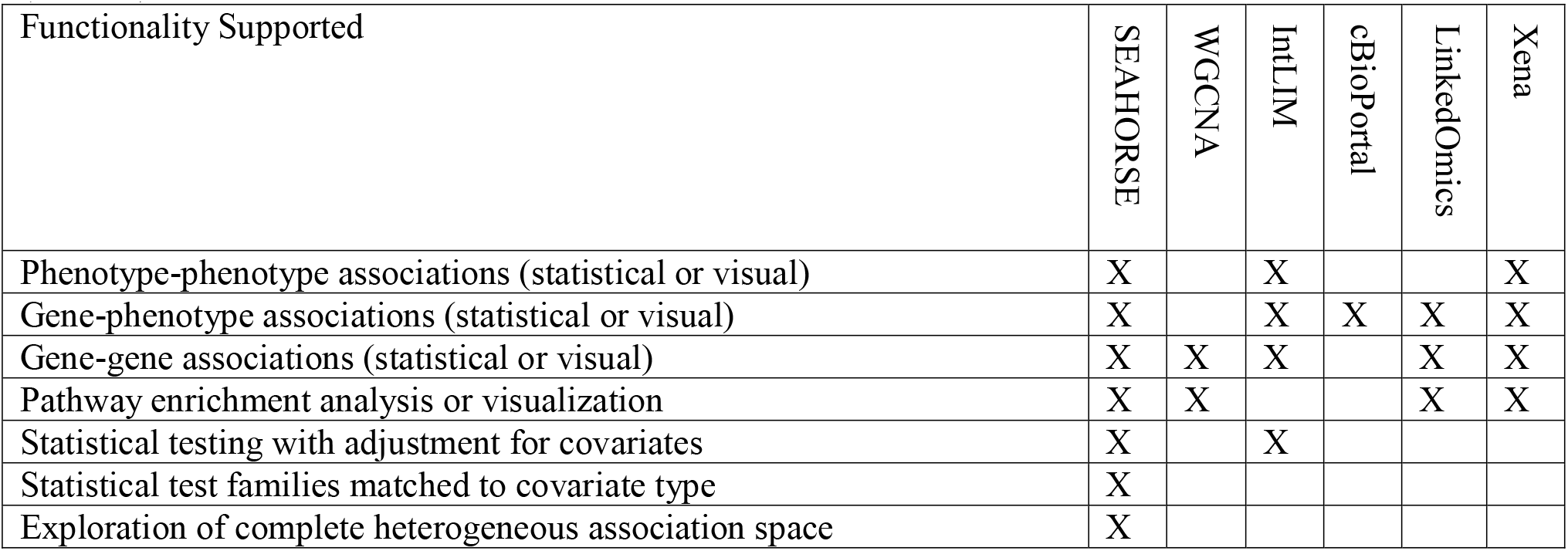
A comparison of the functionalities supported by SEAHORSE and those supported by other tools.

**Table 2.** Statistical tests used for measuring associations between variables within the GTEx data set. The Fisher-Freeman-Halton (FFH) Test is used instead of the Chi-squared test if the number of expected samples in any cell of the expected contingency table is too low to generate reliable results—SEAHORSE uses the *chi*.*sq*() function in the *stats* R package, which sets this threshold at five.

|  | Clinical:<br>Dichotomous | Clinical:<br>Nominal | Clinical:<br>Continuous | Gene<br>Expression |
| --- | --- | --- | --- | --- |
| Clinical:<br>Dichotomous | Chi-squared /<br>Cramer’s V OR<br>FFH | Chi-squared /<br>Cramer’s V OR<br>FFH | Welch’s *t*-test | *limma* moderated<br>*t*-test |
| Clinical:<br>Nominal |  | Chi-squared /<br>Cramer’s V OR<br>FFH | ANOVA | ANOVA |
| Clinical:<br>Continuous |  |  | Spearman<br>Correlation | Spearman<br>Correlation |
| Gene<br>Expression |  |  |  | Pearson<br>Correlation |

The SEAHORSE statistical framework is available as a user-friendly tool within the Network Zoo analytical ecosystem, implemented within the *netZooR* package ^15^ and available from (https://github.com/netZoo/netZooR). We include a tutorial for both navigating the SEAHORSE portal and running the statistical framework on a custom data set in Supplemental File 1. The SEAHORSE portal is available at https://seahorse.networkmedicine.org/.

## Results

### Systematic all-by-all association analysis creates a searchable discovery space

SEAHORSE is designed to support exploratory analysis across multiple classes of variables measured within large data sets.

GTEx v8 release ^2, 16, 17^ has data on 948 individual donors that includes RNA-seq expression from samples representing 43 unique tissues in the body (a reduced subset of the 54 presented in GTEx; see Methods) as well as whole-genome sequencing data from which single-nucleotide polymorphisms (SNPs) can be inferred. GTEx also provides a fairly extensive collection of information about the samples and individuals from whom they were collected, including health-related variables such as height, weight, sex, race, age, cause of death, smoking status, and other relevant information ^18^. The availability of such comprehensive data has already enabled many important analyses, including investigations into phenotypic associations with gene expression and the biological interaction networks that drive transcription ^19–21^, as well as trait association studies that include expression quantitative trait locus (eQTL) analyses ^22–25^. TCGA has collected extensive molecular and phenotypic data from more than 20,000 primary cancer and matched normal samples across 33 organ sites and many annotated cancer types and subtypes ^26^. These data include whole-genome sequencing, whole-transcriptome profiling, and, in many cases, additional data such as miRNA expression, DNA methylation, proteomics, or other data. Many individuals who are represented in TCGA have molecular data on matched tumor and adjacent normal samples, and some have tumors and distant metastases. Associated with each TCGA sample are extensive clinical and phenotypic data on the disease, as well as limited treatment data.

The key design principle behind SEAHORSE is that the system should support not just one type of association, but many, and do so using statistical tests appropriate to the variable types under comparison. That requirement led us to adopt a matrix of test types rather than a single metric of association. Continuous– continuous associations were evaluated by Spearman correlation; dichotomous–continuous associations were assessed using *t*-tests; nominal—continuous associations were assessed using Analysis of Variance (ANOVA); and associations between dichotomous and/or nominal variables were evaluated using contingency-based tests (Table 1). For the Chi-squared Test, the Fisher’s Exact Test, ANOVA, Welch’s *t*-test, and the *limma* moderated *t*-test, all *p*-values were adjusted using FDR correction. Correction was performed with respect to each phenotype within each tissue. The goal was not to force all variables into a uniform representation, but to preserve the heterogeneity of the data while enabling all-by-all interrogation.

The scale of the resulting analysis is substantial. In GTEx, after excluding identifier fields and descriptive comments, we retained 153 phenotypic variables for systematic analysis. Pairwise testing among those variables alone across 43 tissues yielded 341,008 phenotype–phenotype associations. More importantly, testing each phenotype against the expression level of every gene in each tissue produced 125,246,938 phenotype–gene associations. We also computed tissue-specific gene–gene correlations—a total of 10,269,910,410 correlations. These quantities make a simple point: the space of plausible associations in even a single public resource is vastly larger than what any individual investigator would examine by hand.

The decision to precompute these results for exploration in the SEAHORSE portal is therefore not a convenience feature; it is the enabling architecture of the resource. Exploratory science requires short feedback loops. If every question requires re-running a large computational workflow, then only a narrow set of questions will ever be posed. By storing summary statistics, *p*-values, ranked association lists, and downstream enrichment results in indexed data structures, SEAHORSE empowers users to iterate quickly among ideas. One can ask, for example, which phenotypes are associated with age in GTEx; what biological process in each tissue is most strongly correlated with body mass index based on gene expression; or what genes track with the expression of a selected marker in a chosen tissue. The platform is optimized for the phase of analysis that occurs before a formal hypothesis is fully articulated.

### The user interface is designed to promote serendipitous rather than purely targeted discovery

The value of exhaustive computation depends on usability. To that end, the SEAHORSE portal presents its results through a web interface designed explicitly to facilitate unexpected discovery rather than only targeted retrieval. Users begin by selecting the data set of interest, currently GTEx or TCGA. For GTEx, the portal summarizes the cohort through distributions of sampled tissues and key demographic variables, then exposes a query interface that includes phenotype lists, tissue selectors, and gene search boxes. Selecting a phenotype presents its distribution and the corresponding significant associations with other phenotypes or with gene expression. Selecting a gene allows the user to survey phenotype associations and tissue-specific gene–gene associations (Figures 3, 4).

This structure matters scientifically because it encourages lateral movement through the data. An investigator may begin with a variable of obvious interest, but the interface immediately reveals additional associated variables and pathways that were not part of the starting question. We have found this to be one of the most useful aspects of the platform and have used it to generate hypotheses motivating studies published by our group ^27^. In a focused study, one often wants to validate whether a given pathway or phenotype behaves as expected in an external resource. But just as often, exploration reveals that the most interesting signals are the ones that were not initially anticipated. The web interface is therefore not an accessory to the analysis; it is part of the logic of the discovery process.

### Height is associated with multiple disease contexts across human tissues

To illustrate how SEAHORSE can generate unexpected hypotheses, we turned to a deliberately open-ended example: the relationship between height and tissue-specific gene expression in GTEx. Height is an attractive variable for this purpose because it is easily measured, well-studied in epidemiology, and not typically the starting point for tissue-wide transcriptional analysis. We asked, across each of the 43 GTEx tissues, which genes correlate with subject height, then ranked those genes and performed pre-ranked gene set enrichment analysis using KEGG, Gene Ontology (GO) Biological Process, and Reactome annotations (Supplemental Tables S1-S2) ^28–31^. We then examined biological terms that recurred across tissues.

### Associations with recurrent enrichment of cancer-related transcriptional programs

The most striking result from the KEGG analysis was the recurring enrichment of disease-associated pathways among genes correlated with height. Most notably, the KEGG Legacy term “Pathways in Cancer” was significantly associated with height in 17 tissues following FDR adjustment; additionally, pathways representing cancer-associated pathways were significantly associated with height, including GO pathways “Macroautophagy” (24 tissues) and “Regulation of WNT Signaling Pathway” (23 tissues) and Reactome pathways “Translation” (27 tissues) and “DNA Repair” (25 tissues). This observation was surprising to us, not because the epidemiologic literature lacks support for a height–cancer relationship, but because the mechanism linking the two has remained unsettled. Large cohort analyses have reported positive associations between adult height and overall or site-specific cancer incidence ^32^, and a meta-analysis suggested that this is a robust and recurrent observation across many malignancies ^33^. Giovannucci argued that a satisfying mechanistic explanation was lacking, with plausible contributors including increased cell number, growth factor biology, and metabolic factors—but failed to identify an unifying transcriptional model ^33^.

SEAHORSE does not by itself establish causality, nor does it determine whether the observed enrichment reflects intrinsic tissue programs, developmental history, endocrine signaling, cell composition, or latent confounding. But it does provide an important clue: across many tissues, height is associated not simply with isolated genes but with coordinated pathway-level changes involving canonical cancer biology. That pattern is exactly the sort of observation that a discovery engine should expose. It takes an epidemiologic association often discussed at the population level and points to a plausible molecular mechanism at the individual level that can be examined more rigorously in independent studies.

The scope of the gene expression signal we found is especially notable. A signal confined to one or two tissues might be dismissed as noise or an idiosyncratic, tissue-specific process. A signal recurring in the majority of tissues suggests that height may be linked to systemic biological processes with implications for cell growth, signaling, or tissue organization. That does not mean all tissues are affected identically, or that pathway membership alone implies directional equivalence across cancers. But the repeated appearance of cancer-related transcriptional programs across such a large fraction of the body’s tissues argues that anthropometric variation and tissue-level molecular state may be more tightly coupled than is generally appreciated.

This example also illustrates why all-by-all analyses matter. It is unlikely that a conventional targeted analysis of GTEx would have begun by asking whether height is associated with pathway-level shifts in cancer biology across many tissues. Height is often treated as a covariate, not a discovery variable. Yet once the full association space is made searchable, such questions become natural to ask—and, critically, the most interesting answers may emerge only after the fact. In that sense, the height result captures the central logic of SEAHORSE: exhaustive comparison creates opportunities for serendipity that targeted analysis would rarely produce.

### Height is also associated with recurrent neurologic, immune and cardiovascular disease-related pathway signals

The height example yielded more than one class of unexpected pathway association; we found, immune, cardiovascular, and neurologic pathways to be enriched across multiple tissues. The most commonly enriched immune pathways included the GO Biological Processes “Wound Healing” (26 tissues) and “Defense Response to Virus” (20 tissues) and the Reactome pathways “Neutrophil Degranulation” (25 tissues), “Signaling by the B Cell Receptor” (26 tissues), “Influenza Infection” (25 tissues), “Dengue Virus Infection” (22 tissues), and “HIV Infection” (20 tissues). A few studies have investigated the relationship between height and immune response, with mixed results^34, 35^. However, because immune response is a crucial process in the context of many diseases, the associations discovered using SEAHORSE present a serendipitous area for further study, with implications for individualized medicine based on anthropometric variation.

Among the most common KEGG Legacy terms associated with height was “cardiac muscle contraction,” which was significant in 16 tissues. Also prominent, appearing in 14 tissues, was “arrhythmogenic right ventricular cardiomyopathy (ARVC),” a genetic heart condition characterized by the replacement of normal heart muscle with fatty or fibrous tissue in the right ventricle, which can lead to arrhythmias and potentially life-threatening complications. Vascular pathways included Reactome pathway “Cell Surface Interactions at the Vascular Wall” (21 tissues), pertaining specifically to the interaction of platelets and leukocytes with the endothelium via adhesion molecules (R-HSA-202733), and “Regulation of Vasculature Development” (18 tissues). These findings are interesting because the clinical literature on height and cardiovascular disease is mixed but far from empty. Reviews have noted associations between stature and multiple cardiac phenotypes, with one of the strongest and most reproducible links involving atrial fibrillation and venous thromboembolism. Large-scale epidemiologic and Mendelian randomization analyses have suggested that greater height may increase the risk of atrial fibrillation even as it shows inverse or more complex relationships with some vascular outcomes^36–39^, and venous thromboembolism is known to be positively associated with increased height in both men and women^40–43^.

Here again, SEAHORSE does not solve the mechanism, but it points toward one. The recurrent association between height and cardiovascular transcriptional programs across many tissues suggests that anthropometric variation may capture molecular processes relevant to structural or electrophysiologic cardiovascular phenotypes. Moreover, SEAHORSE suggests that the adhesion molecules in platelet-leukocyte interactions present an interesting area for future research in understanding the relationship between height and venous thromboembolism. Finally, the recurrent association between height and immune response merits further investigation, particularly in the context of other factors such as socioeconomic status and exposure to pathogens in early life, which can impact both height and immune response^44–46^.We were particularly struck by the appearance of ARVC as, unlike atrial fibrillation, this is not a height association that is well established in the literature. That discrepancy is exactly what makes the result interesting. It may reflect pathway overlap with broader cardiac remodeling processes, or it may suggest a new line of investigation into shared molecular architecture among height-associated cardiac phenotypes.

Several additional disease-related enrichments associated with height were also consistent with published findings. Shorter stature has been linked in some studies on Parkinson’s disease risk (KEGG Legacy pathway “Parkinson’s Disease,” enriched in 21 tissues), possibly through decreased density of neurons in the substantia nigra pars compacta ^47^ (although studies report the opposite association with height ^48^), and a body of work connects shorter height or height loss to cognitive decline, dementia, or Alzheimer’s disease ^49–53^ (KEGG Legacy pathway “Alzheimer’s Disease,” enriched in 19 tissues). We do not claim that SEAHORSE resolves questions surrounding these published findings. Rather, the point is that a discovery system should recover a mixture of expected, literature-supported, and genuinely surprising associations. That is what we observe here.

### Height-disease associations persist when adjusting for covariates

While SEAHORSE is primarily designed for hypothesis generation using all-by-all associations, *netZooR* also supports follow-up linear regression analyses on gene-phenotype data to adjust for covariates. To demonstrate how such follow-up analyses can further inform our understanding of the possible relationship between height and tissue-specific pathways, we evaluated the associations between gene expression and height in each tissue, adjusting for body mass index (BMI), sex, and age (Equation 1). We calculated *p*-values were using the moderated *t*-statistic for *β*_1_, and the moderated *t*-statistic for _1_ was used as the input score for GSEA.

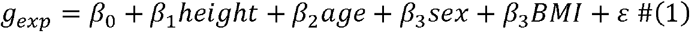

We found that the aforementioned pathways corresponding to cancer pathogenesis, cardiovascular modeling, immune response, and neurodegenerative disease also persisted after adjusting for these covariates (Table 3).

**Table 3.** Commonly enriched cancer, immune, cardiovascular, and neurological pathways and the number of tissues in which they are enriched in the hypothesis generation model and the follow-up linear model with adjustment for age, sex, and BMI.

| Pathway | Number of Tissues in Which Enriched |  |
| --- | --- | --- |
|  | Unadjusted | Adjusted |
| Translation | 27 | 26 |
| Wound Healing | 26 | 26 |
| Signaling by the B Cell Receptor | 26 | 26 |
| Neutrophil Degranulation | 25 | 25 |
| Influenza Infection | 25 | 25 |
| DNA Repair | 25 | 24 |
| Macroautophagy | 24 | 24 |
| Regulation of WNT Signaling | 23 | 23 |
| Dengue Virus Infection | 22 | 22 |
| Cell Surface Interactions at the Vascular Wall | 21 | 22 |
| Parkinson’s Disease | 21 | 20 |
| HIV Infection | 20 | 21 |
| Defense Response to Virus | 20 | 20 |
| Alzheimer’s Disease | 19 | 20 |
| Regulation of Vasculature Development | 18 | 18 |
| Pathways in Cancer | 17 | 18 |
| Cardiac Muscle Contraction | 16 | 16 |
| ARVC | 14 | 15 |

The persistence of these pathway enrichment results after adjustment suggests that the associations between height and cancer, immune, cardiovascular, and neurological pathways are not due to indirect effects of age, sex, or BMI. While this result still does not imply causality or define the mechanism underlying these associations, it suggests that treating height as a discovery variable in the context of these conditions may be beneficial for understanding disease heterogeneity. Tissue-specific results and full enrichment counts are available from Supplemental Tables S3-S4.

### Across TCGA, age is recurrently associated with canonical oncogenic and matrix-remodeling programs

We next asked whether the same exploratory strategy could reveal cross-cancer transcriptional patterns in TCGA. Cancer is often described as a disease of aging, largely because the incidence of many adult cancers increases sharply with age. Yet the relationship between age and cancer phenotype is not simple. For several tumor types, younger patients present with more advanced disease or biologically aggressive subtypes, while older patients often experience worse outcomes because of both tumor-intrinsic and host-related factors ^54–56^. Age, therefore, offers an informative axis along which to search for recurrent tumor transcriptional programs.

Using harmonized RNA-seq profiles and phenotype data for 33 TCGA cancer types provided by recount3 ^57^, we computed tumor-type-specific associations between gene expression and clinical variables using the same general statistical framework applied to GTEx. We then ranked genes by their association with patient age and performed pathway enrichment analysis on the significant genes in each comparison (Supplemental Tables S5-S6). Across tumor types, age was repeatedly associated with pathways involving WNT signaling, translation, cell differentiation, and cell cycle mechanisms, each of which was identified in more than 15 cancers. More broadly, the aggregate KEGG term “Pathways in Cancer” was significantly associated with age in 17 tumor types. The roles played by several of these pathways in cancer pathogenesis with respect to increasing age are supported by previous work. The Reactome pathway “mRNA Splicing” was significant in 20 tumor types, consistent with the role of age-associated aberrant alternative splicing in cancer ^58^, The GO pathway “Mitotic Spindle Checkpoint” was significant in 19 tumor types, consistent with the role of aging-related spindle checkpoint failures in cancer-associated aneuploidy. ^59^ Finally, the Reactome pathway “Telomere Maintenance” was significant in 16 tumor types – this pathway regulates shortening of telomeres, which characterizes aging and is known to impact cancer development.^60^

These findings are not merely unsurprising confirmations that cancer pathways matter in cancer. Their value lies in recurrence and organization. Age is not a tumor-specific mutation or a histopathologic subtype; it is a host attribute. The fact that age-associated transcriptional signatures repeatedly converge on WNT signaling, translation, cell differentiation, and cell cycle mechanisms suggests that aging may help shape a conserved set of tumor-associated programs across tissue contexts. This is consistent with the broader idea that changes in the aged microenvironment, in systemic signaling, or in the biology of aging tissues may influence how tumors initiate, progress, or evade control.

From a translational perspective, the result also suggests that age-associated programs may merit closer attention in biomarker development or therapeutic stratification. If age is associated with recurring pathway-level changes across tumor types, then some of the heterogeneity currently attributed solely to tissue of origin or mutational state may also reflect age-dependent transcriptional context. In this sense, SEAHORSE does not replace focused mechanistic studies, but it helps identify recurrent themes that may be worthy of further pursuit.

### SEAHORSE recovers positive control associations in GTEx and TCGA

To evaluate recovery of expected (“positive control”) associations between genes and phenotypes using SEAHORSE, we measured the significance of the following well-known associations in both GTEx and TCGA:

- Cell cycle gene expression level correlation (pairwise)
- Mitosis gene expression level correlation (pairwise)
- Ribosomal gene expression level correlation (pairwise)
- Sex and Y-chromosome gene expression level association
- Sex and height association

When averaging across tissues, we found that all cell cycle, mitosis, and ribosomal gene correlations were above 0.5 in GTEx (a total of 1,582,034,245 GTEx gene correlations were above 0.5, placing the positive controls in the top 15.4^th^ percentile) and 0.75 in TCGA (a total of 370,695,278 TCGA gene correlations were above 0.75, placing the positive controls in the top 3.1^st^ percentile) (Supplemental Figure S1). The lowest correlation values in GTEx involved cell cycle gene *CCNB1* and included aorta, gastroesophageal junction, esophagus muscularis, atrial appendage, and pituitary gland. These low correlations may reflect biological differences in the respective tissues; for instance, coordination of cell cycle genes in both cardiomyocytes in the atrial appendage and vascular smooth muscle cells in the aorta may differ from that of other tissues because both are long-lived cells ^61, 62^. Abnormal regulation of *CCNB1* in cardiovascular tissue is also linked to restenosis (blockage of a coronary artery previously treated with a stent) – because cardiovascular disease was reported in 19.2% of GTEx donors (188) and cardiovascular conditions including cardiovascular disease, cardiac failure, cerebrovascular disease, heart attack, and stroke were listed as a first immediate cause of death for 45.2% of GTEx donors (443), it is possible that a considerable percentage of donors exhibit abnormal *CCNB1* expression in cardiovascular tissue ^63^.

Associations between sex and Y-chromosome gene expression level and between sex and height were highly significant in both data sets (Supplemental Figure S2). Detail regarding selection of genes and variables is detailed in Supplemental Table S7 along with tissue-specific results. We conclude from this result that SEAHORSE recovers known biological associations between genes and phenotypes.

### SEAHORSE scales to large cohort studies on a laptop

To evaluate the scalability of SEAHORSE to large cohorts in a laptop environment, we ran SEAHORSE on TCGA and GTEx data using a MacBook Pro laptop running macOS 14.7.6 with 16 GB RAM and an Apple M3 chip with 8 cores. Obtaining results for gene-gene correlations and for gene-phenotype and phenotype-phenotype associations for all GTEx tissues required a total of 18.49 hours with an average increase in usage of 1.35 GB RAM. Obtaining these results for all TCGA tissues required a total of 16.86 hours with an average increase in usage of 1.57 GB RAM (Supplemental Table S8).

## Discussion

The principal contribution of SEAHORSE is not simply that it stores precomputed results, but that it changes how investigators can interact with large cohort data sets. Most public molecular resources are optimized either for data distribution or for relatively narrow forms of query. Although those are essential functions, they do not fully exploit the fact that such data sets contain a vast, mostly unsearched association space. By computing, indexing, and visualizing that space, SEAHORSE turns the data set itself into a hypothesis generator.

We want to emphasize that this is a complement to, not a replacement for, conventional hypothesis-driven science. Discovery engines are most valuable when they produce observations that can be tested through orthogonal analysis, independent data, or direct experimentation. Indeed, one useful feature of SEAHORSE is that it can serve two distinct purposes: it can generate unexpected hypotheses and provide a convenient means to validate whether associations identified elsewhere are present in GTEx or TCGA. In that respect, the platform sits at the interface between exploration and confirmation.

The height example provides perhaps the clearest demonstration of this logic. Height is not typically considered a central driver of tissue-wide transcriptional organization, yet SEAHORSE identifies recurring pathway-level associations that align with epidemiologic observations in cancer and cardiovascular disease. We do not interpret these findings as final explanations. Rather, they are precisely the sort of structured, biologically coherent observations that should trigger deeper, directed studies. The fact that the signal emerges across many tissues further suggests that anthropometric variables may encode broad system-level biology that is underappreciated in standard transcriptomic analysis.

At the same time, the platform necessarily raises issues of multiplicity and interpretation. The sheer number of associations tested means that false positives are unavoidable if one naively interprets every nominally significant relationship as a discovery. We view this not as a flaw specific to SEAHORSE, but as an inherent feature of large-scale exploratory analysis. SEAHORSE is intended to surface candidate observations whose plausibility can then be assessed through recurrence across tissues, pathway-level coherence, literature triangulation, external data sets, or direct experiments. In other words, the output of a discovery engine is not a conclusion; it is a prioritized, often unexpected, starting point.

Several limitations also point toward future development. The current release emphasizes RNA-seq and phenotype associations and, for TCGA, focuses on primary tumor samples. Yet both GTEx and TCGA, and many other modern resources, contain additional data types including mutation, methylation, copy number, proteomics, and imaging-derived variables. Extending SEAHORSE across these modalities is an obvious and important next step. The same is true for expansion to new cohorts, including resources such as Trans-Omics for Precision Medicine (TOPMed), All of Us, the Million Veterans Program, the American Association for Cancer Research (AACR) Genomics Evidence Neoplasia Information Exchange (GENIE), and the United Kingdom (UK) Biobank, as well as the large collection of recount3-standardized transcriptomic studies. We also see value in extending the resource to include inferred gene regulatory networks and other association models ^22, 64–68^ as an additional data type for interrogation; our recent development of a robust, streamlined Nextflow pipeline for network inference makes this extension to large-scale data sets feasible ^69^. As the diversity of measured and biologically constrained inferred variables increases, the value of a unified framework for heterogeneous all-by-all discovery only grows.

More broadly, we think the argument made by SEAHORSE is methodological as much as biological. Biology now possesses many data sets whose scale exceeds the question-formulation bandwidth of individual investigators. All scientific discoveries begin with observations that can be more deeply studied and verified or falsified. In the era of big data, discovery requires infrastructure. If we want to find the “unknown unknowns,” we must build systems that make them searchable. SEAHORSE is one such system. Its purpose is to make large public data sets more useful not only for answering the questions we already know how to ask, but for helping us discover better questions.

## Methods

### GTEx data set and phenotype processing

As a first implementation of SEAHORSE, we used data from GTEx version 8 (accessed October 17, 2021) ^1^, selecting the 948 genotyped individuals for whom both RNA-seq data and clinical phenotype information were available. RNA-seq values were downloaded from the GTEx portal as transcripts per million (TPM) and used without additional normalization in the resource-generation workflow. The GTEx phenotype tables, obtained through dbGaP study accession phs000424.v8.p2, contained 189 variables, which we categorized as nominal, dichotomous, or continuous.

We preprocessed these data by replacing values of “Not Reported,” “Unknown,” “Not Performed,” or “Indeterminate” with NA values, converting temporal variables reported in HH:MM format or “H hour(s), M minute(s)” format to minutes, and consolidating data where appropriate to remove redundant categories. We then removed variables corresponding to identifiers or clinical notes as well as invariant and redundant variables. After filtering, we retained 153 phenotype variables for association analysis, including 16 library metadata variables such as refrigeration time and ischemic time for the sample. A full record of modifications to the phenotypic data is available in Supplementary Table S9.

The tissues included in the current release represent a reduced subset of the full GTEx tissue panel and correspond to the 43 tissues for which the implemented workflow generated stable summaries and association catalogs. Within each tissue type, we removed genes that had counts of <1 TPM in at least 5% of samples. We removed two samples that were designated as “biological outliers” in the GTEx portal (https://gtexportal.org/); that is, those samples from donors with a large chromosomal duplication or deletion or who have undergone gender-affirming surgery (as described in https://gtexportal.org/home/faq). We consolidated brain tissue samples into three separate groups due to expression heterogeneity described by Paulson et al ^70^: Brain-0 (hypothalamus, amygdala, anterior cingulate cortex, hippocampus, cortex, substantia nigra, spinal cord (cervical c-1), and frontal cortex), Brain-1 (cerebellum, cerebellar hemisphere), and Brain-2 (caudate (basal ganglia), nucleus accumbens (basal ganglia), and putamen (basal ganglia)). Finally, we log-transformed the TPM gene expression data (Equation 1).

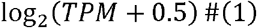

### Statistical framework for GTEx association analysis

All statistics were computed using SEAHORSE as implemented in *netZooR* by calling the *seahorse*() function. We computed phenotype–phenotype and phenotype–gene associations and gene–gene correlations. For phenotype–phenotype analysis, all pairwise associations among the retained variables were evaluated, with statistics being computable for 341,008 associations. The statistical test used depended on variable type. Continuous–continuous associations were assessed with Spearman correlation; dichotomous or nominal categorical variables were tested against continuous variables using *t*-tests or ANOVA; and categorical–categorical associations were evaluated with contingency-based approaches. The objective was to apply an appropriate statistical model to each pairwise association rather than force all variables into a single metric. For phenotype–gene associations, each phenotype was tested against the expression of each gene in each tissue using Spearman correlation, t-tests, or ANOVA as appropriate, producing 125,246,938 associations. For gene–gene analysis, within each tissue we calculated Pearson correlations across genes, producing 10,269,910,410 correlations. We selected Spearman correlation rather than Pearson correlation for phenotype data because it is more appropriate for data with a wide variety of distributions, including nonnormally distributed data or data with outliers, while Pearson correlation offers runtime advantages and has been shown to outperform Spearman correlation in normalized gene expression data from medium-sized cohorts, including GTEx ^71–74^. Input data and resulting statistics were stored in indexed R data frames keyed by the variables involved in each comparison.

### Pathway enrichment analysis

To move from long ranked lists of genes to interpretable biological summaries, we performed pathway enrichment on phenotype-associated genes using pre-ranked gene set enrichment analysis (GSEA) implemented through the *fgsea* R package. For continuous phenotypes, ranks were determined by correlation, with both positive and negative enrichment reported. For nominal phenotypes, ranks were determined by −log_10_ p, where *p* is the *p*-value of the ANOVA, and only positive enrichment was reported. For dichotomous phenotypes, ranks were determined by *t-*statistic, with both positive and negative enrichment reported. Enrichment analyses were performed against KEGG, Gene Ontology Biological Process, and Reactome gene sets ^28, 75, 76^, with analysis limited to pathways with between 15 and 500 genes. Pathway files were downloaded from the Molecular Signatures Database (MSigDB) through https://www.gsea-msigdb.org/gsea/msigdb/collections.jsp^77^. For compatibility with the gene expression data, we mapped all gene symbols to Ensembl IDs using the *mapIds*() function from the *AnnotationDbi* R package, allowing for multiple values. For the examples emphasized in this paper, including the analyses of height in GTEx and age in TCGA, we ranked significant terms (FDR-adjusted *p*-value < 0.05) by the number of tissues or tumor types in which they appeared as significant and then examined the recurrent patterns among those terms. Complete enrichment results for selected analyses are available through the SEAHORSE portal.

### TCGA data set and processing

SEAHORSE also includes exploratory analyses using The Cancer Genome Atlas (accessed November 21, 2021), which contains molecular and phenotype information across more than 20,000 tumor and matched normal samples from 33 cancer types. The current implementation focuses on primary tumor RNA-seq and phenotype data. Gene expression and phenotype data for the 33 cancer types were downloaded from recount3, a uniformly processed resource for large-scale RNA-seq studies ^57^. We transformed the raw count data to TPM using the *transform_counts*() function from *recount3* and log-scaled the TPM values. Similarly to GTEx, we removed genes with counts of <1 TPM in at least 5% of samples and limited our analysis to samples that included both phenotypic data and expression data. Moreover, we limited our analysis to primary tumors only. We computed tumor-type-specific associations using the SEAHORSE functionality in the netZooR package in the same manner as for GTEx. Pathway enrichment analysis was then carried out using the same gene sets used for GTEx analysis using the SEAHORSE functionality.

Phenotypic data were preprocessed similarly to GTEx. Timestamps were converted to Unix time values, redundant groups were consolidated, and identifier data, file names, file md5sums, clinical notes, invariant variables, and inconsistently formatted variables were removed. After filtering, we retained 709 phenotype variables for association analysis, including 147 library metadata variables such as source site, Formalin-Fixed Paraffin-Embedded (FFPE) processing, batch number, read count, days between index date and date of sample receipt, and record update timestamps. See Supplementary Table S10 for descriptions of phenotype variable transformations.

### User interface and implementation

The SEAHORSE portal is exposed as a web-based query tool that allows users to browse precomputed summary statistics, association tables, and visual summaries. The portal includes data-source selection, phenotype menus, tissue selectors, and gene query boxes. For GTEx, selecting a phenotype displays its empirical distribution together with significantly associated phenotypes and tissue-specific gene associations. For both GTEx and TCGA, users can inspect ranked results and, where relevant, follow those results into downstream pathway summaries. The code used to compute the association catalog was written in R and is distributed under a Massachusetts Institute of Technology (MIT) license through the project’s public repository, which is hosted on GitHub (https://github.com/netZoo/netZooR). The association catalog is stored in AWS Simple Storage Service (S3) buckets, and the backend is implemented using AWS Lambda functions integrated with AWS Application Programming Interface (API) Gateway to expose those functions through Hypertext Transfer Protocol (HTTP) endpoints. API rate and burst limits are set to 5 requests per second (RPS) and 15 RPS, respectively, for all queries. To enhance reproducibility, an online Jupyter notebook titled “Uncovering Associations among Genes and Phenotypes with SEAHORSE” is provided through Netbooks ^78^ (https://netbooks.networkmedicine.org), enabling users to reproduce core portions of the workflow.

### Illustrative analyses

The example analyses described in this manuscript were selected to demonstrate the kinds of observations that emerge when the entire association space is made searchable. For height in GTEx, we computed tissue-specific associations between subject height and gene expression, ranked genes by significance, and then performed pre-ranked GSEA. For age in TCGA, we computed cancer-type-specific associations between patient age and gene expression and performed the same enrichment workflow. These examples were not chosen because they were the only associations observed, but because they illustrate the platform’s central use case: identifying coherent, potentially important biological patterns that might not be obvious from conventional, narrowly targeted analysis.

## Data availability

SEAHORSE is available as a web-based query tool at https://seahorse.networkmedicine.org/. A reproducible notebook implementation of core analyses is available through Netbooks (https://netbooks.networkmedicine.org) as “Uncovering Associations among Genes and Phenotypes with SEAHORSE,” and a user guide is available in Supplemental File 1. Full results and each tissue and tumor type are available from Zenodo, along with full inputs for each tumor type, originally obtained through recount3 (https://rna.recount.bio/). Inputs for GTEx are not provided for download because the GTEx phenotype data are controlled access—these are accessible through dbGaP study accession phs000424.v8.p2 upon request.

Links to the records are listed below:

- TCGA inputs: https://zenodo.org/records/21300598
- GTEx results (part 1): https://zenodo.org/records/21417165
- GTEx results (part 2): https://zenodo.org/records/21431005
- GTEx results (part 3): https://zenodo.org/records/21463147
- GTEx results (part 4): https://zenodo.org/records/21480298
- TCGA results (part 1): https://zenodo.org/records/21302428
- TCGA results (part 2): https://zenodo.org/records/21328475
- TCGA results (part 3): https://zenodo.org/records/21347083
- TCGA results (part 4): https://zenodo.org/records/21364228

GTEx phenotype data were obtained from dbGaP study accession phs000424.v8.p2, and TCGA RNA-seq summaries were obtained through recount3 (https://rna.recount.bio/).

## Code availability

Software used to calculate associations and generate the current SEAHORSE resource was written in R and is distributed under the MIT License through the project repository associated with the manuscript (https://github.com/netZoo/netZooR/tree/master). The code used to reproduce the results presented in this manuscript is available from https://github.com/QuackenbushLab/seahorse-networks. Front-end and back-end code for the SEAHORSE portal are available from https://github.com/QuackenbushLab/seahorse-frontend and https://github.com/QuackenbushLab/seahorse-backend, respectively.

## Acknowledgements

ES, TE, KHS, and JQ were supported by National Institutes of Health grants R35CA220523 and R01HG011393; DD and JQ were supported by U24CA231846. The project was also supported by Amazon Web Services through the Harvard Data Science Institute.

