## Supplementary Materials for "SEAHORSE: A Serendipity Engine Assaying Heterogeneous Omics-Related Sampling Experiments"

SEAHORSE User Guide

SEAHORSE combines two separate tools for serendipitous discovery: the *netZooR* R package function *seahorse*(), which implements the statistical framework and can be applied to custom data sets, and the SEAHORSE portal, an online user interface for exploring associations in the TCGA and GTEx data sets.

### Navigating and Interpreting SEAHORSE Portal

#### SEAHORSE Portal Field Dictionary

The SEAHORSE portal draws data from .TSV.GZ files which are stored in AWS S3 buckets for GTEx and TCGA, respectively. Each file is read in as a separate table and concatenated across GTEx and TCGA.

##### ensembl2symbol

| **Field** | **Type** | **Description** | **Example** |
| --- | --- | --- | --- |
| *alias* | Character | Alias for the gene (duplicate of symbol) | MYC |
| *ensembl_id* | Character | Ensembl identifier for the gene | ENSG00000136997 |
| *symbol* | Character | HGNC-approved gene symbol | MYC |
| *entrez_id* | Integer | Entrez identifier for the gene | 4609 |
| *dataset_id* | Character | Dataset from which this entry is sourced | gtex |

##### gene_expression

| **Field** | **Type** | **Description** | **Example** |
| --- | --- | --- | --- |
| *ensembl_id* | Character | Ensembl identifier for the gene | ENSG00000136997 |
| *gtex_id* | Character | Identifier for the sample | phv00174086.v7.p2.c1 |
| *level* | Real | Gene expression level (logTPM) | 1.25 |
| *dataset_id* | Character | Dataset from which this entry is sourced | gtex |

##### expression_correlation

| **Field** | **Type** | **Description** | **Example** |
| --- | --- | --- | --- |
| *gene_a* | Character | Ensembl identifier for the first gene in the pair | ENSG00000136997 |
| *gene_b* | Character | Ensembl identifier for the second gene in the pair | ENSG00000000001 |
| *tissue* | Character | Tissue in which the correlation was calcualted | whole_blood |
| *correlation* | Real | Correlation between gene_a and gene_b | 0.7 |
| *dataset_id* | Character | Dataset from which this entry is sourced | gtex |

##### gsea

| **Field** | **Type** | **Description** | **Example** |
| --- | --- | --- | --- |
| *pathway* | Character | Name of pathway | REACTOME_CELL_CYCLE |
| *pvalue* | Real | Enrichment p-value | 0.005 |
| *padj* | Real | BH-adjusted enrichment p-value | 0.05 |
| *lod2err* | Real | Precision of p-value estimation | 1.27 |
| *es* | Real | Enrichment score | -0.65 |
| *nes* | Real | Normalized enrichment score | -3.47 |
| *size* | Integer | Size of pathway | 97 |
| *ranks* | Character | Vector of ranks of genes in pathway | {1,2,50} |
| *leading_edge* | Character | Vector of genes driving enrichment | {ENSG00000138796,  ENSG00000181092} |
| *varname* | Character | Name of variable | HGHT |
| *tissue* | Character | Tissue in which enrichment was evaluated | whole_blood |
| *dataset_id* | Character | Dataset from which this entry is sourced | gtex |

##### metadata

| **Field** | **Type** | **Description** | **Example** |
| --- | --- | --- | --- |
| *gtex_id* | Character | Identifier for the sample | phv00174086.v7.p2.c1 |
| *tissue* | Character | Tissue in which enrichment was evaluated | whole_blood |
| *varname* | Character | Name of variable | HGHT |
| *value* | Text | Value of variable | 171 |
| *dataset_id* | Character | Dataset from which this entry is sourced | gtex |

##### metadata2expression

| **Field** | **Type** | **Description** | **Example** |
| --- | --- | --- | --- |
| *varname* | Character | Name of variable | DTHCOD |
| *ensembl_id* | Character | Ensembl identifier for the gene | ENSG00000136997 |
| *tissue* | Character | Tissue in which enrichment was evaluated | whole_blood |
| *test* | Character | Type of test performed | ANOVA |
| *test_statistic* | Real | Test statistic (correlation or nominal p-value) | 0.005 |
| *pvalue* | Double precision | Adjusted p-value | 0.05 |
| *dataset_id* | Character | Dataset from which this entry is sourced | gtex |

##### metadata2metadata

| **Field** | **Type** | **Description** | **Example** |
| --- | --- | --- | --- |
| *category_a* | Character | Name of first variable | DTHCOD |
| *category_b* | Character | Name of second variable | DTHFUCOD |
| *tissue* | Character | Tissue in which enrichment was evaluated | whole_blood |
| *test* | Character | Type of test performed | ANOVA |
| *test_statistic* | Real | Test statistic (correlation or nominal p-value) | 0.005 |
| *pvalue* | Double precision | Adjusted p-value | 0.05 |
| *dataset_id* | Character | Dataset from which this entry is sourced | gtex |

#### Getting Started – Overview of Associations in GTEx and TCGA

When using the SEAHORSE Portal, users must first know which tissue type and set of phenotypes, library metadata variables, and genes they wish to explore. Once users have made these selections, they may use the SEAHORSE Portal to explore associations between variables of interest.

We provide tables of the most significant gene-gene, gene-phenotype, phenotype-phenotype, and phenotype-pathway associations within each GTEx tissue and TCGA tumor type to give users a starting point for meaningful exploration of the Portal using very stringent cutoffs (correlation > 0.99 and FDR-adjusted p-value < 5e-20) (Supplementary Tables S11-S12). Users can also generate larger tables on their own using the *writeSigTable*() function in netZooR on the GTEx and TCGA results, which are downloadable from Zenodo.

#### Navigating Cohort, Tissue, and Phenotype

After navigating to the SEAHORSE Portal (https://seahorse.networkmedicine.org/#/), users may select either GTEx or TCGA to explore the respective cohorts. The portal does not currently provide a programmatic API. Data are accessed through the graphical web interface, where users specify tissues, genes, phenotypes, and other search parameters using interactive controls.

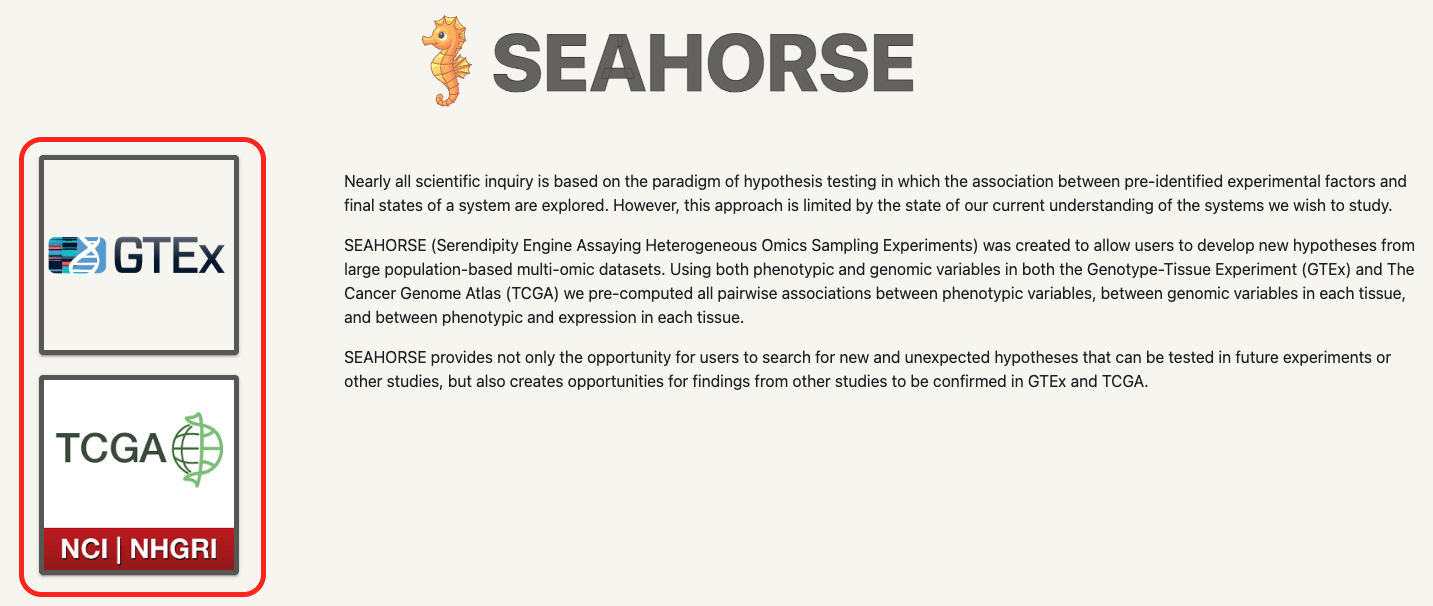

After selecting a database, users may then navigate between tissue / tumor types using a drop-down window at the top of the screen.

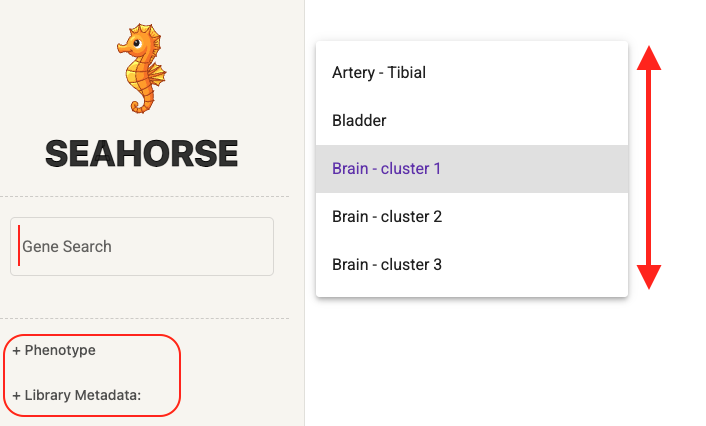

Within the tissue / tumor type, users may select phenotypes or library metadata variables from the side panel to visualize a phenotype / library metadata summary histogram and phenotype-phenotype, phenotype-gene, and phenotype-pathway associations.

Users may also enter genes into the text box to visualize a gene expression summary histogram and phenotype-gene and gene-gene associations.

At any time, users may return to the home page by clicking on the SEAHORSE icon in the upper-left corner of the screen.

#### Summary Histogram – Phenotype, Library Metadata, or Gene

For the selected phenotype, library metadata variable, or gene, the histogram shows the distribution of that phenotype. This plot appears at the top of the page. The example below is the histogram for a continuous phenotype.

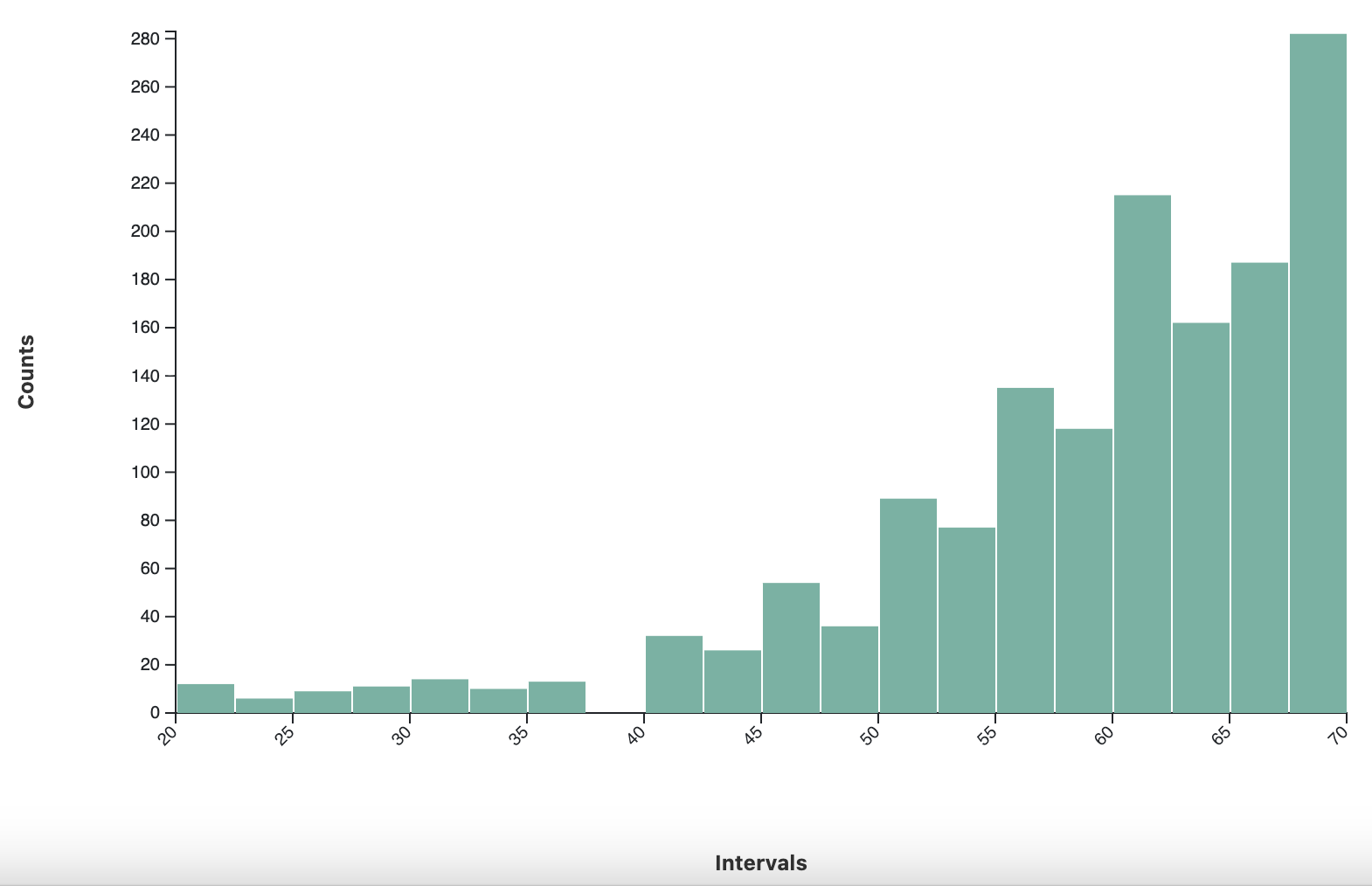

#### Phenotype-Phenotype, Phenotype-Metadata, Phenotype-Gene, and Gene-Gene Associations

Users may view associations as an interactive table or a series of plots in the following ways:

- **Phenotype-Phenotype:**
  - Select a phenotype from the side panel 🡪 Expand the “Phenotype” section
- **Phenotype-Metadata:**
  - Select a library metadata variable from the side panel 🡪 Expand the “Phenotype” section, OR
  - Select a phenotype from the side panel 🡪 Expand the “Library Metadata” section
- **Phenotype-Gene:**
  - Select a phenotype from the side panel 🡪 Expand the “Gene” section, OR
  - Search for a gene in the text box 🡪 Expand the “Phenotype” section
- **Phenotype-Pathway:**
  - Select a phenotype from the side panel 🡪 Expand the “Pathways” section
- **Metadata-Metadata:**
  - Select a library metadata variable from the side panel 🡪 Expand the “Library Metadata” section
- **Metadata-Gene:**
  - Select a library metadata variable from the side panel 🡪 Expand the “Phenotype” section, OR
  - Search for a gene in the text box 🡪 Expand the “Library Metadata” section
- **Metadata-Pathway:**
  - Select a library metadata variable from the side panel 🡪 Expand the “Pathways” section
- **Gene-Gene:**
  - Search for a gene in the text box 🡪 Expand the “Gene” section

##### Heatmap, Box Plot, or Scatterplot

For the selected phenotype, library metadata variable, or gene, plots are used to visualize associations with top six most significantly associated phenotypes, library metadata variables, or genes. Users may select each plot to show a zoomed-in visualization of that plot.

When a nominal or dichotomous variable is compared against another nominal or dichotomous variable, the associations are visualized as a heatmap, as shown in the example below.

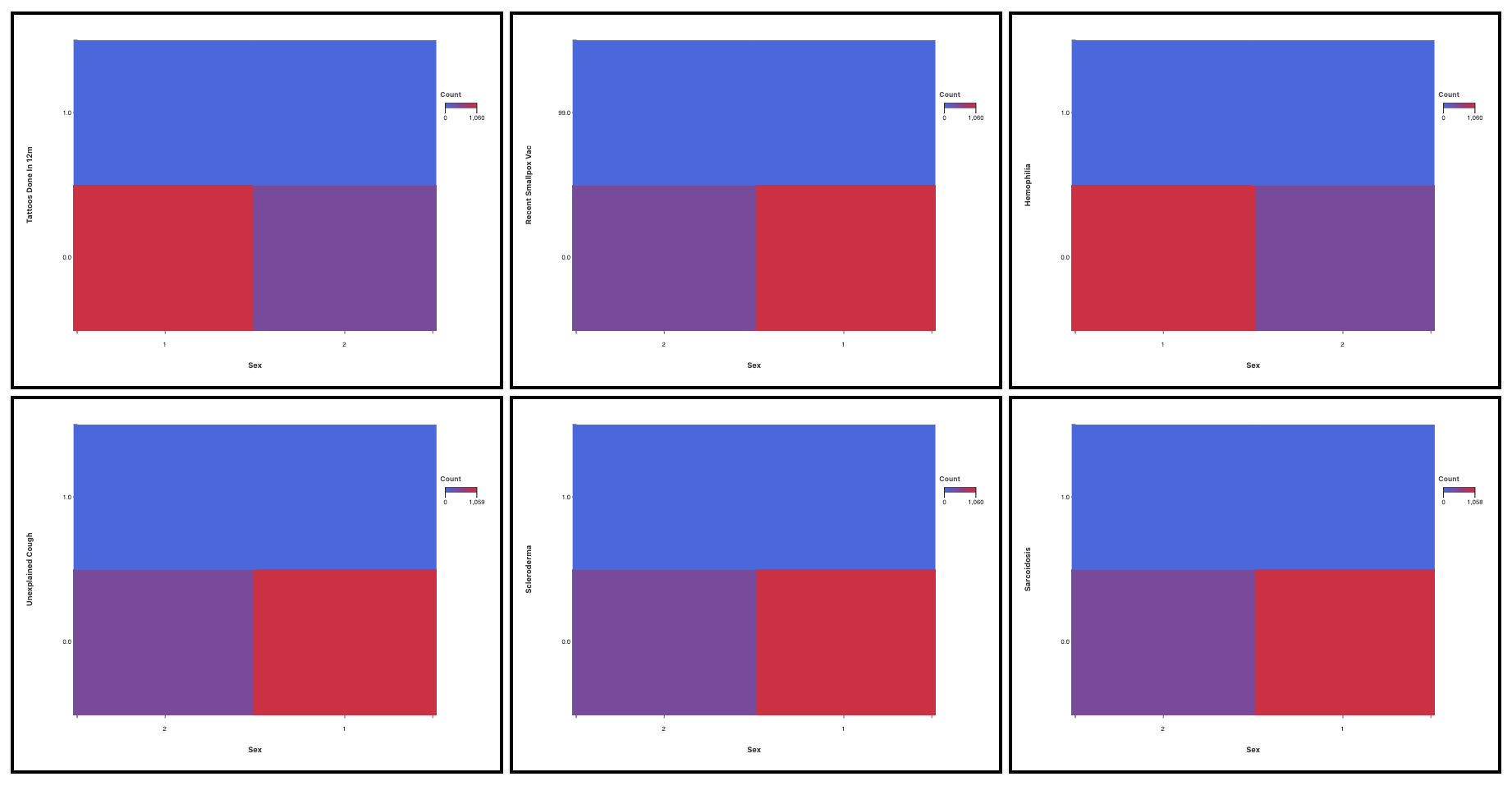

When a continuous variable is compared against a dichotomous or nominal variable, a box plot is shown (example below).

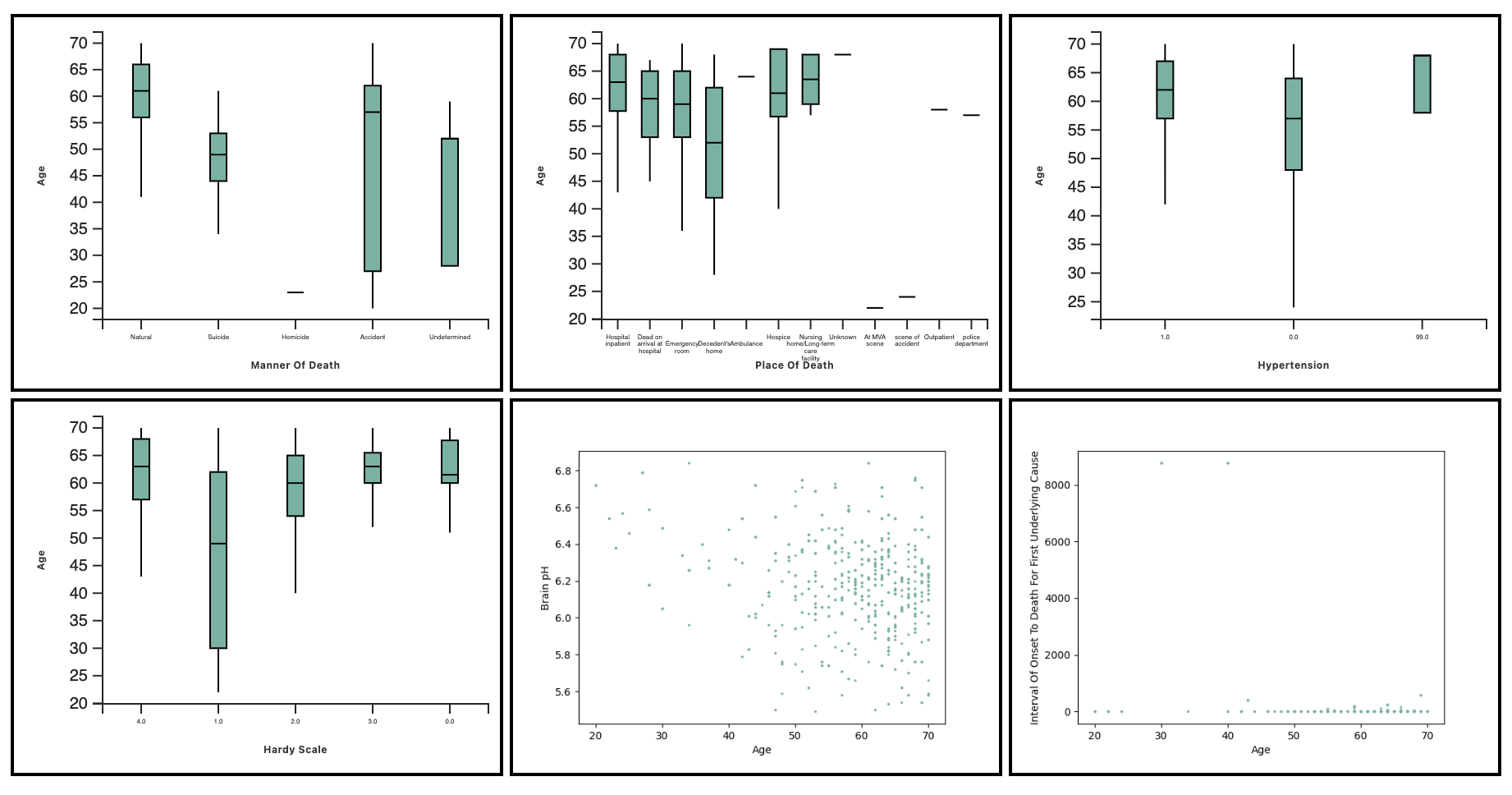

When a continuous phenotype is compared against another continuous phenotype, a scatterplot is shown.

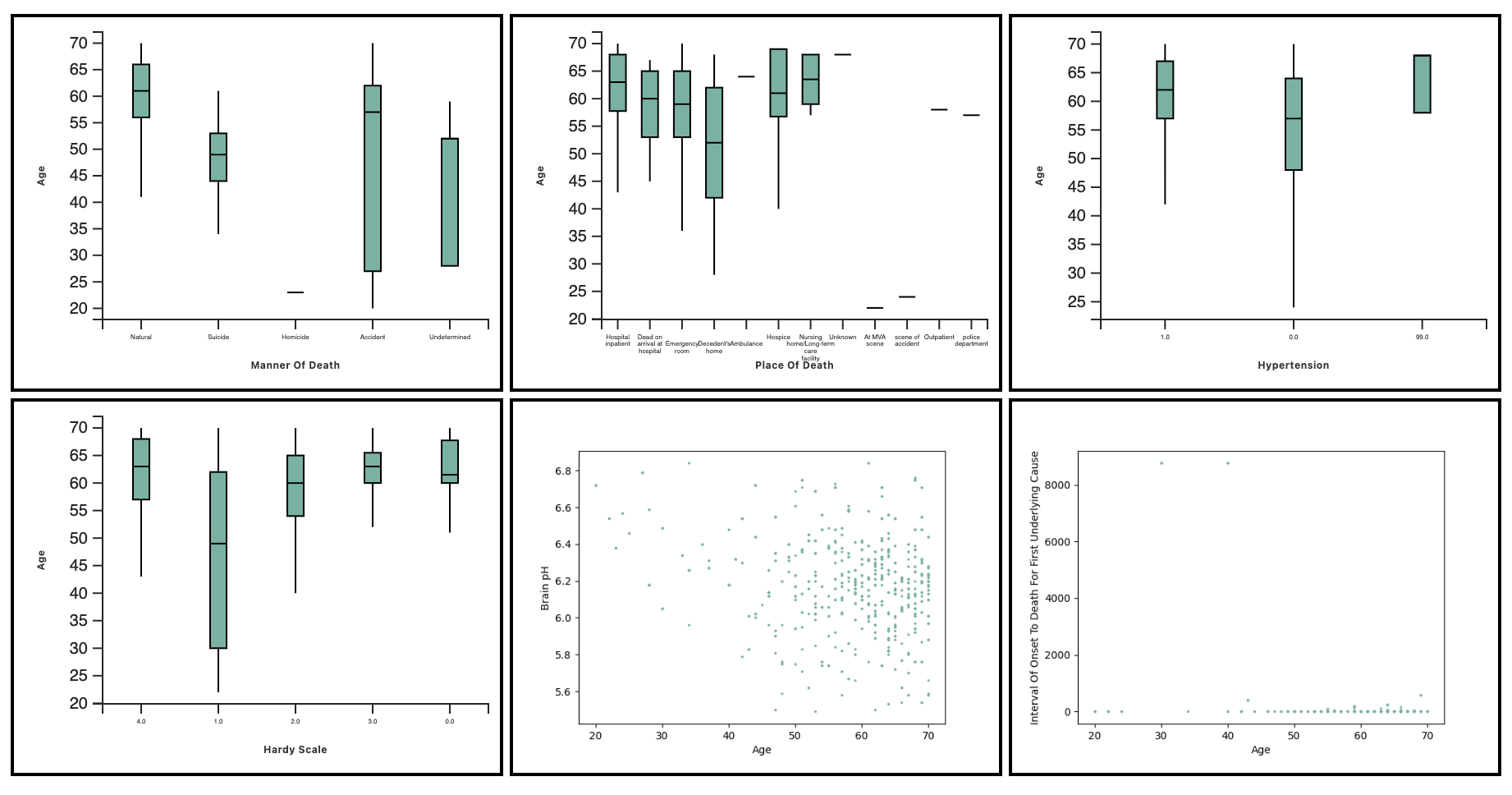

##### Interactive Table

A table is also shown directly below the plot that details test statistics (correlations or *p*-values) and adjusted *p*-values for all associations. Users may use the arrows below the table to navigate to the variable of interest, sorted by association significance. Users may also click on links in the first column of the table to generate a heatmap, boxplot, or scatterplot in real time.

#### Phenotype-Pathway and Metadata-Pathway Associations

Users may expand the Pathways Section to view rug plots for the pathways most significantly associated with the selected phenotype or library metadata variable. In the rug plot for each pathway, the black bars represent the rankings of test statistics for each gene within the pathway. In general, the more significant a pathway is, the more black bars are expected to concentrate at the left tail of the distribution in comparison to the right tail.

Selecting the pathway name will open the MSigDB entry for that pathway in a separate tab. Selecting the “Tissues” button next to the pathway will display a list of other tissues in which the pathway is significant. An example of the rug plot distribution is shown below.

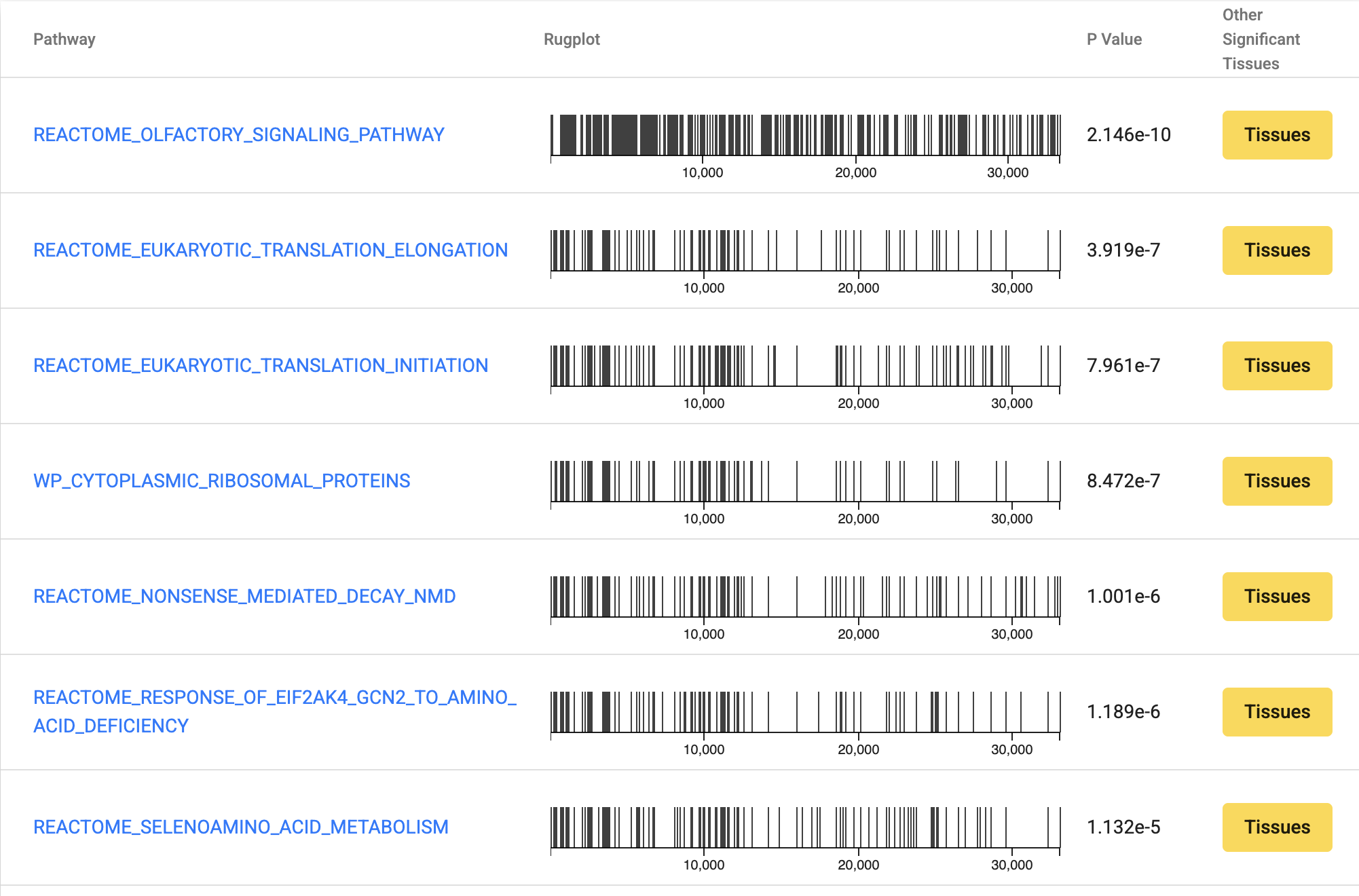

### Running the SEAHORSE Statistical Framework on Your Data

The following instructions describe how to run SEAHORSE on a custom data set using the *seahorse*() function in the *netZooR* R package. Note that, in the R package, library metadata are treated as a subset of the phenotypic data. Therefore, when the term “phenotype” is used here, library metadata variables are also included.

1. Preprocess your data for SEAHORSE analysis. You must ensure that:
   a. The sample labels match between your phenotype and expression data.
   b. Expression data are scaled appropriately for use with *limma*. This means normalized counts that have been log-scaled or transformed using *voom* or *DESeq*.
   c. In your expression data, the rows are genes and the columns are samples.
   d. In your phenotype data, the rows are samples and the columns are phenotypes.
   e. You have created a phenotype dictionary. This is a vector of values corresponding to variable type for each phenotype – e.g., for a phenotype matrix containing the columns “case / control”, “age”, and “tumor grade”, the corresponding dictionary would contain “dichotomous”, “continuous”, and “nominal”, in that order.
   f. You have loaded a pathway file containing a list of genes for each pathway (e.g., a GMT format file).
2. Install netZooR: <https://github.com/netZoo/netZooR>
3. Run SEAHORSE in R using the following command. If you are running SEAHORSE on multiple tissue types or cohorts, you will need to run it separately for each.

| **seahorse()** | **Computes and returns all associations of interest for a single data set** |
| --- | --- |
| expression | The preprocessed expression data |
| phenotype | The preprocessed phenotype data |
| phenotype_dictionary | The phenotype dictionary |
| pathways | The pathway data |
| compute_gene_cor | Whether or not to compute the correlations between genes (TRUE / FALSE). Default is TRUE. |
| compute_phenotype_cor | Whether or not to compute the correlations between phenotypes. Default is TRUE. |
| compute_gene_phenotype_cor | Whether or not to compute the correlations between genes and phenotypes and their corresponding GSEA analysis (TRUE / FALSE). Default is TRUE. |
| assoc_method | Method to use for correlation analysis on phenotype-phenotype and phenotype-gene pairs. Can be “pearson”, “kendall”, “spearman”, or “linear” (a linear regression model with adjustment for covariates). Default is “spearman”. If using “linear”, set compute_phenotype_cor = FALSE, as this method only applies to gene-phenotype relationships. |
| pval_adj_method | Method for p-value adjustment. All valid inputs to the *p.adjust*() method in the *stats* R package are valid. Default is “none” (no adjustment). |
| usage_report_file | Path to the RDS file where the usage report (memory and runtime analysis) will be stored. If NULL (default), the report will not be saved. |
| verbose | Whether or not to print statements notifying the user of run status (TRUE / FALSE). Default is “FALSE”. |

### Visualizing Your Data Using the SEAHORSE R Package

The SEAHORSE R package includes visualizations analogous to the SEAHORSE Portal, allowing users to perform similar exploration of their own data sets. Documentation for each visualization function is given in the table below, where the function is described along with its input arguments. The term “phenotype” as used here includes both phenotypic variables and library metadata variables.

| **rugPlot()** | **Generates a rug plot for a single pathway-phenotype association** |
| --- | --- |
| result | The SEAHORSE result |
| pathwayName | The pathway of interest |
| phenotypeName | The phenotype of interest |
| pathways | The pathway data, formatted as input to SEAHORSE |
| **summaryHistogramGene()** | **Generates a summary histogram for a gene** |
| expression | The gene expression data, formatted as input to SEAHORSE |
| geneName | The gene of interest |
| breaks | The number of breaks in the histogram |
| **summaryHistogramPhenotype()** | **Generates a summary histogram for a phenotype** |
| phenotype | The phenotype data, formatted as for SEAHORSE |
| phenotypeName | The phenotype of interest |
| phenotype_dictionary | The phenotype dictionary, formatted as inupt to SEAHORSE |
| breaks | The number of breaks in the histogram |
| **plotAssociation()** | **Plots the association between two variables (genes or phenotypes)** |
| expression | The gene expression data, formatted as input to SEAHORSE |
| phenotype | The phenotype data, formatted as input to SEAHORSE |
| phenotype_dictionary | The phenotype dictionary, formatted as input to SEAHORSE |
| variableX | The variable of interest to plot on the x-axis (a phenotype or gene) |
| variableY | The variable of interest to plot on the y-axis (a phenotype or gene) |
| variableTypeX | Either "gene" or "phenotype" (x-axis variable) |
| variableTypeY | Either "gene" or "phenotype" (y-axis variable) |
| **writeTable()** | **Write the table containing the results (gene-gene, gene-phenotype, phenotype-pathway, or phenotype-phenotype) for one phenotype or gene.** |
| result | The full SEAHORSE result |
| phenotype | The phenotype data, formatted as input to SEAHORSE |
| dictionary | The phenotype dictionary, formatted as input to SEAHORSE |
| pathways | The pathway data, formatted as input to SEAHORSE |
| variable | The variable of interest by which to filter (a phenotype or gene) |
| variableType | Either "gene" or "phenotype" |
| resultType | The name of the result component you wish to save as a table ("coexpression", "phenotype_association", "phenocor", or "GSEA") |
| tmpFile | The temporary file you wish to create for constructing the table |
| resultFile | The location where you wish to save the table |

### FAQ

**Q: I am getting a “Package ‘x’ is required but not installed” error. What does this mean?**

A: Several of the SEAHORSE functions in NetZooR require specialized R packages that are not automatically installed when the R package is built. If you see “Package ‘x’ is required but not installed”, install the R package on your machine and try to run SEAHORSE again.

**Q: I am getting one of the following error messages. How can I fix these?
“Phenotype ‘x’ set to nominal but has 2 levels or fewer”**

**“Phenotype ‘x’ set to dichotomous but has more than 2 levels”**

**“Phenotype ‘x’ set to continuous but cannot be converted to numeric”**

A: When you generate the phenotype dictionary, it is important to make sure that the values in the dictionary match what is in the data and that the order of the dictionary matches the order of the columns in the phenotype file. If a variable has character data with two levels or fewer, it should be listed as “dichotomous” in the dictionary. If it has more than two levels, it should be listed as “nominal”. Only numeric variables should be listed as “continuous”.

**Q: Why am I seeing an error that the file I specified for the usage report “could not be created”?**

A: SEAHORSE tests that the usage report file can be created by opening the file, writing a small amount of data to it, and then deleting it. If you receive this error message, you may have specified an invalid path or have insufficient permissions to create or edit the file.

**Q: Why am I seeing the warning “Could not compute empirical Bayes statistics for this phenotypic variable. Returning empty list.”?**

A: For dichotomous phenotypes, SEAHORSE performs a *limma*-moderated t-test, and for nominal phenotypes, SEAHORSE performs an ANOVA. In both cases, it runs the *lmFit*() and *eBayes*() functions from the *limma* R package on all genes in parallel with respect to the phenotype. A common scenario causing *eBayes*() to fail is when the vast majority of samples have missing data for this phenotype. In this scenario, SEAHORSE will return NA values for both the phenotype-gene and phenotype-pathway associations.

**Q: Why am I seeing the warning “Phenotype has only one level. Returning NA for all gene associations.”?**

A: This occurs when a phenotype takes only one value across all samples, so associations with that phenotype cannot be computed. In this scenario, SEAHORSE will return NA values for both the phenotype-gene and phenotype-pathway associations. Although retaining these phenotypes won’t cause SEAHORSE to fail, you can decrease runtime by removing phenotypes with one level before running SEAHORSE.

**Q: Why am I seeing the warning “Less than 2 nonzero marginals in the contingency table - Chi-square will return NA”?**

A: This occurs when phenotype-phenotype associations are being computed, both phenotypes are either dichotomous our nominal, and the contingency table used to compute a Chi-square or FFH test has less than 2 nonzero row marginals or column marginals. This occurs due to the distribution of samples across both phenotypes. In this scenario, the association cannot be computed, and SEAHORSE will return NA values for the phenotype-phenotype association statistics.

**Q: Why am I seeing the warning “In this phenotype pair, all continuous values are missing for all but one phenotype level - NA result will be returned”?**

A: This occurs when phenotype-phenotype associations are being computed, one phenotype is nominal, the other phenotype is continuous, and values of the continuous phenotype are missing for all but one level of the nominal phenotype (e.g., number of cigarettes smoked per day may be missing for donors who did not smoke, making the association between cigarettes smoked per day and smoking status meaningless). In this scenario, the association cannot be computed using ANOVA, and SEAHORSE will return NA values for the phenotype-phenotype association statistics.

**Q: Why am I seeing the warning “Some phenotypes did not have sufficient sample sizes to perform a t-test - NAs will be returned”?**

A: This is essentially the previous warning (“In this phenotype pair, all continuous values are missing for all but one phenotype level - NA result will be returned”) but for dichotomous phenotypes instead of nominal phenotypes. SEAHORSE handles these cases in the same way as it does for the ANOVA.

**Q: Why am I seeing the warning “Could not compute t-test (it is possible that variance is too low). Returning NA.”?**

A: This occurs when phenotype-phenotype associations are being computed, one phenotype is dichotomous, the other phenotype is continuous, and the *t*-test cannot be computed. As the warning states, a common scenario in which this occurs is when the variance of the continuous phenotype is very low across one or both levels of the dichotomous phenotype. In this scenario, the association cannot be computed using a *t*-test, and SEAHORSE will return NA values for the phenotype-phenotype association statistics.

**Q: Why are some of my phenotype-phenotype associations being computed using the Chi-square test and others using the FFH test?**

A: SEAHORSE defaults to using the Chi-square test to find associations across two dichotomous or nominal phenotypes because it is less computationally intensive than the Fisher-Freeman-Halton (FFH) test. However, the Chi-square test is known to produce unstable results in some conditions, such as when one of the expected counts in the contingency table is less than five. In these scenarios, SEAHORSE does not report the unstable Chi-square result and instead reports the FFH test result.

**Q: Why are Cramer’s V values not returned for all phenotype-phenotype associations?**

A: Cramer’s V can only be computed using the results from a Chi-square test. In scenarios where an FFH test, ANOVA, or *t*-test is used instead, Cramer’s V values will be set to NA.
