## Supplementary Figures for "SEAHORSE: A Serendipity Engine Assaying Heterogeneous Omics-Related Sampling Experiments"

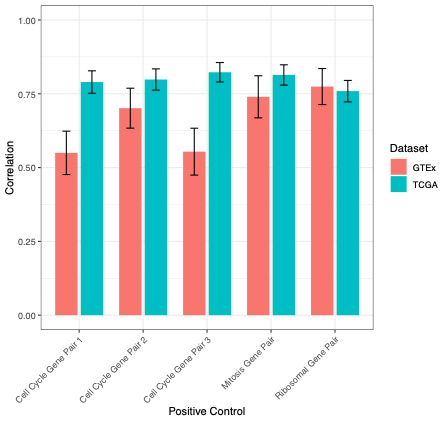

Supplemental Figure S1. Mean correlation values resulting from SEAHORSE analysis across GTEx tissues (pink) and TCGA tumor types (blue) for each positive control gene pair, with 95% confidence intervals.

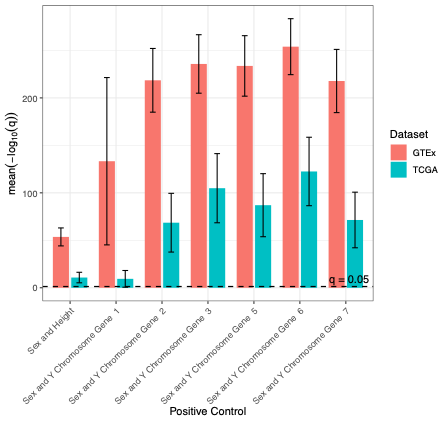

Supplemental Figure S2. Mean *q*-values (FDR-adjusted *p*-values) resulting from SEAHORSE analysis across GTEx tissues (pink) and TCGA tumor types (blue) for each positive control gene-phenotype or phenotype-phenotype pair, with 95% confidence intervals.
